# A regulatory protein that represses sporulation in *Clostridioides difficile*

**DOI:** 10.1101/2020.02.25.964569

**Authors:** Diogo Martins, Aristides L. Mendes, Jessica Antunes, Adriano O. Henriques, Mónica Serrano

## Abstract

Bacteria that reside in the gastrointestinal tract of healthy humans are essential for our health, sustenance and well-being. About 50 to 60% of those bacteria have the ability to produce resilient spores, important for the life cycle in the gut and for host-to-host transmission. A genomic signature for sporulation in the human intestine was recently described, which spans both commensals and pathogens such as *Clostridioides difficile*, and contains several genes of unknown function. We report on the characterization of a signature gene, *csiA*, which, as we show, is involved in the control of sporulation initiation in *C. difficile*. Spo0A is the main regulatory protein controlling entry into sporulation and we show that an in-frame deletion of *csiA* results in increased sporulation, and increased expression of *spo0A* per cell. Spo0A also drives transcription of the *spoIIA* and *spoIIG* operons, coding for the first forespore-(σ^F^) and mother cell-specific (σ^E^) RNA polymerase sigma factors. Strikingly, deletion of *csiA* increases expression of the *spoIIG* operon, but not that of the *spoIIA* operon. Increased expression of *spoIIG* results in increased production and proteolytic activation of pro-σ^E^, suggesting that normally, the levels of active σ^E^ are limiting for sporulation. While other regulatory proteins affect both sporulation and several processes during the transition phase of growth, including toxin production or motility, deletion of the *csiA* gene does not alter the expression of the genes coding for the TcdA and TcdB cytotoxins or the genes involved in motility. Thus, our results establish that CsiA acts to modulate sporulation by reducing expression of the *spo0A* gene.

## Introduction

Many members of the Firmicutes phylum produce endospores (hereafter spores for simplicity), resilient and metabolically dormant structures that can remain viable for long periods of time. A large proportion of recently identified members of the human gut microbiota are spore formers and many of which are anaerobic organisms (Almeida *et al*., 2019, Browne *et al*., 2016). For anaerobic bacteria, pathogens or commensals, the production of oxygen-resistant spores is an important mechanism for transmission between hosts and persistence inside and outside the host (Browne *et al*., 2016). Gut spore forming bacteria are important members of the human commensal microbiota that only in rare cases cause opportunistic infections (Atarashi *et al*., 2011, Browne *et al*., 2016). Other spore formers, however, have evolved to become dedicated pathogens that can cause a striking variety of diseases. Despite variations in disease presentation, the infectious agent is often the spore, with toxins produced by the actively growing cells that result from spore germination playing a central role in the pathophysiology of infection (Zhu *et al*., 2018). One example is *Clostridioides difficile*, a spore-former obligate anaerobe. *C. difficile* is a major nosocomial pathogen and the main causative agent of a range of intestinal diseases associated with antibiotic therapy in adults (Carroll & Bartlett, 2011).

Soon after the initiation of sporulation, the sporulating cell undergoes an asymmetric division originating two compartments of unequal sizes, the large mother cell and the small forespore (Errington, 2003, Higgins & Dworkin, 2012, Piggot & Hilbert, 2004)(Fig. 1A). The forespore and the mother cell will follow different lines of gene expression (Fimlaid *et al*., 2013, Pereira *et al*., 2013, Saujet *et al*., 2013). Entry into sporulation involves phosphorylation of the Spo0A response regulator, a transcription factor conserved among spore formers and essential for sporulation initiation in all organisms in which its function was experimentally assessed (Abecasis *et al*., 2013, Galperin *et al*., 2012, Traag *et al*., 2013). The role of Spo0A in sporulation has been studied in detail in *Bacillus subtilis* (reviewed in (Deakin *et al*., 2012, Fujita & Losick, 2005, Molle *et al*., 2003, Pettit *et al*., 2014, Sonenshein, 2000)). In pre-divisional cells, the Spo0A-encoding gene, *spo0A*, is transcribed under the control of σ and σ (Pettit *et al*., 2014).

**Figure 1.**
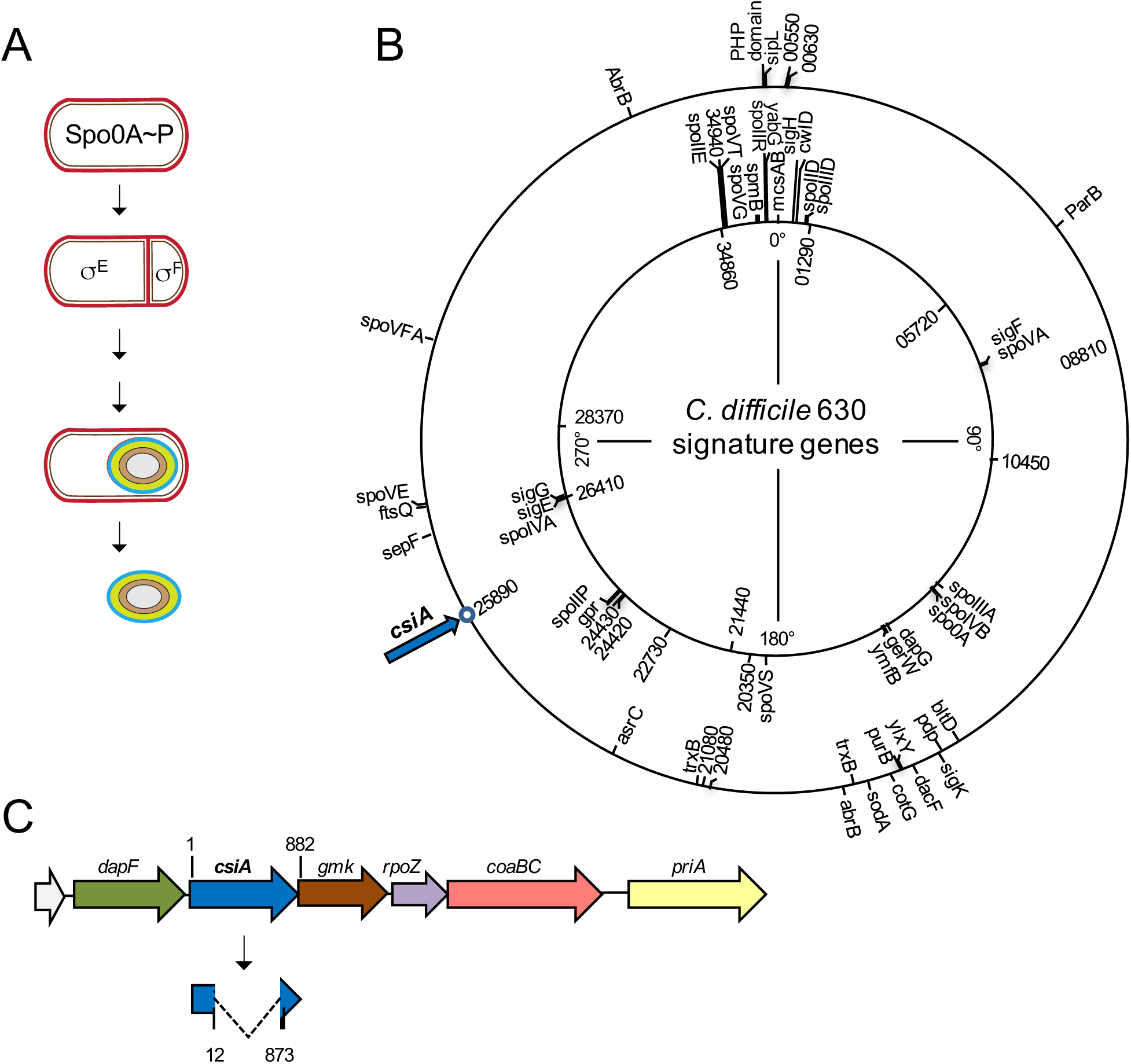
Genomic signature of sporulation. **A:** main stages of sporulation starting with asymmetric division of the rod-shaped cell and the production of the larger mother cell and the smaller forespore (the developing spore) through the development of full spore refractivity, until the release of the spore into the environment. The onset of sporulation is controlled by phosphorylation of Spo0A. After asymmetric division, the forespore and the mother cell compartments follow different lines of gene expression. **B:** The inner circle contains the signature defined as those genes present in 90% of sporulating bacteria and in no more than 10% of the remaining bacterial species (Abecasis *et al*., 2013). The outer circle has the remaining genes identified in a genomic signature of sporulation within the human intestinal microbiome (Browne *et al*., 2016). Both signatures are enriched with known sporulation-associated genes involved in spore morphogenesis and germination. Genes not specifically associated with a sporulation stage or uncharacterized genes are shown inside the circles. The positions of the genes are shown in degrees in the *C. difficile* 630 chromosome. The *csiA* gene (CD25890) is a signature gene not previously characterized (blue arrow). **C:** schematic representation of the *csiA* (CD25890) region of the *C. difficile* 630 Δ*erm* chromosome. The Δ*csiA* in-frame deletion removed codons 5 through 291 of the 293-codons-long *csiA* gene.

Spo0A is post-translationally activated *via* phosphorylation by a phosphorelay consisting of several sensor kinases and phosphortransfer proteins, with multiple points of control of the flow of phosphate to Spo0A. The kinases respond to external signals by auto-phosphorylating and subsequently transferring a phosphate residue *via* the intermediary proteins Spo0F and Spo0B to Spo0A (Jiang *et al*., 2000). In contrast, no Spo0F or Spo0B homologs are found in *C. difficile*, and therefore presumably no phosphorelay is present; rather, Spo0A is believed to be phosphorylated directly by orphan histidine kinases that respond to as yet unknown signals (Steiner *et al*., 2011, Underwood *et al*., 2009, Worner *et al*., 2006). The *C. difficile* genome codes for five orphan kinases, three of which, CD1492, CD1579 and CD2492, share a high degree of sequence similarity with the sensor histidine kinases involved in the *B. subtilis* phosphorelay (Underwood *et al*., 2009). These three kinases were initially proposed to be responsible for Spo0A phosphorylation. CD1579, however, was the only kinase shown to directly phosphorylate Spo0A. CD2492 was proposed to positively regulate sporulation (Underwood *et al*., 2009), however no complementation was shown for the mutant. In contrast, CD1492 is now known to inhibit sporulation initiation and to affect toxin and motility through the regulatory proteins RstA and SigD (Childress *et al*., 2016).

In *C. difficile*, Spo0A controls approximately 300 genes (Fimlaid *et al*., 2013, Pettit *et al*., 2014), which have been linked to biofilm formation, swimming motility, toxin production and sporulation (Dawson *et al*., 2012, Fujita & Losick, 2005, Pettit *et al*., 2014, Underwood *et al*., 2009). Among the latter, are the genes coding for the first cell type-specific regulators of sporulation, (σ^F^ and σ^E^), as well as genes required for the asymmetric division of the cell at the onset of sporulation (Errington, 2003, Higgins & Dworkin, 2012, Pettit *et al*., 2014, Piggot & Hilbert, 2004, Rosenbusch *et al*., 2012).

Following asymmetric division, the mother cell initiates engulfment of the forespore, a process with similarities to phagocytosis (Fig. 1A). At this stage, the forespore is surrounded by an inner and outer membranes, that derive from the polar division septum. Following engulfment completion, two types of peptidoglycan are layered between the inner and outer membranes surrounding the forespore. The surface of the inner spore membrane is the site of assembly of a thin layer of peptidoglycan called primordial cell wall, similar in composition to the vegetative cell wall. A second, thicker layer of chemically distinct peptidoglycan, called the spore cortex, is assembled across the forespore outer membrane, which faces the mother cell cytoplasm. The spore cortex is essential for the attainment and maintenance of the dehydrated state of the spore core, and hence for heat resistance and dormancy (Henriques & Moran, 2007). Finally, in *C. difficile*, the ordered assembly of proteins produced in the mother cell, form the coat and the exosporium (Henriques & Moran, 2007, Paredes-Sabja *et al*., 2014, Stewart, 2015). These proteinaceous layers protect the cortex against the action of peptidoglycan-breaking enzymes, confer protection against small noxious molecules and UV light, and influence the interaction of spores with molecules able to trigger germination (Driks & Eichenberger, 2016, Henriques & Moran, 2007, McKenney *et al*., 2013). In addition, the exosporium plays an important role not only in the adhesion of spores to abiotic surfaces but also to host cells (Stewart, 2015). At the end of the sporulation process the mother-cell lyses, releasing the spore into the environment.

We previously established a genomic signature for sporulation, defined as the genes found in 90% of the endospore forming bacteria, and present in no more than 10% of the non-spore-formers ((Abecasis *et al*., 2013); Fig. 1B). This signature not only allows prediction of spore formation by organisms for which sporulation has not been shown in the laboratory, but it also identified new genes involved in sporulation (Abecasis *et al*., 2013). More recently, a genomic signature for endosporulation in the human gastro-intestinal tract was established ((Browne *et al*., 2016); Fig. 1B). This signature includes 65 genes and is dominated by genes with a known function in sporulation. Approximately 30% of the signature genes, however, have no known function in sporulation and/or code for products with no similarity to known proteins; nevertheless, their presence in a genomic signature for sporulation suggests a role in spore development, at least in the gut. One of these genes is *CD25890* (Fig. 1B) and here we show that deletion of *CD25890* increases the expression of *spo0A*, and also the expression of the Spo0A- and σ^A^-dependent *spoIIG* operon, coding for the first mother cell specific RNA polymerase sigma factor, σ^E^. In contrast, deletion of *CD25890* does not increases the expression of the Spo0A- and σ^H^-dependent *spoIIA* operon, coding for the first forespore-specific sigma factor, σ^F^. Expression of the *spoIIG* operon results in the production of an inactive form of σ^E^, pro-σ^E^. Cleavage of pro-σ^E^ requires a forespore signalling protein, which in *C. difficile* is mostly but not completely dependent on σ (Fimlaid *et al*., 2013, Pereira *et al*., 2013, Saujet *et al*., 2013). Strikingly, increased expression of *spoIIG* in the *CD25890* mutant results in increased production and proteolytic activation of pro- σ^E^, suggesting that the levels of active σ, and not forespore signalling, are a limiting factor for sporulation. Deletion of the *CD25890* gene does not alter the expression of the gene coding for the TcdA cytotoxin, or the genes involved in motility, or biofilm formation. Based on these phenotypes we renamed the *CD25890* locus *csiA*, to reflect its role in the <u>c</u>ontrol of <u>s</u>porulation initiation.

## Results

### Functional analysis of a *csiA* mutant

Phylogenetically, the *csiA* gene and its genomic context are highly conserved within the Firmicutes (Fig. S1). In each genome we examined, CsiA is encoded immediately upstream of the *gmk* gene encoding a guanylate kinase or of the *remA* gene encoding a biofilm regulatory protein (Winkelman *et al*., 2009, Winkelman *et al*., 2013). These genes are always upstream of the *rpoZ* gene, encoding for the RNA polymerase omega subunit. Although its genomic context is conserved, there is no evidence for the possible function of *csiA*. Neverthelss, its inclusion in a genomic signature for sporulation suggested an important role in this developmental process. We started by the phenotypic characterization of a mutant in the *csiA* gene. An in-frame deletion mutant of *csiA,* lacking codons 5 – 291 of the 294-codons-long open reading frame was generated by allelic-coupled exchange (ACE) in the background of the widely used 630Δ*erm* strain (Fig. 1C; Fig. S2; Table S1; (Ng *et al*., 2013)). The mutant was then complemented in single copy upon restoration of the *pyrE* gene (Fig. S2).

**Table 1.**
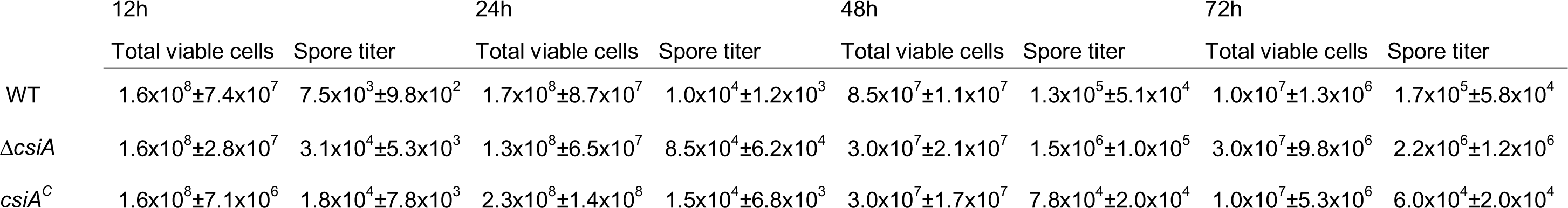
Sporulation efficiency of the ⊗csiA mutant. The total viable cell count and the spore titer was determined 12, 24, 48 and 72 hours following inoculation into sporulation medium. The heat resistant spore count by determining the CFU/ml before and after incubation at 70°C for 10 min. The results shown are averages and standard deviations for three biological replicates.

|  | 12h |  | 24h |  | 48h |  | 72h |  |
| --- | --- | --- | --- | --- | --- | --- | --- | --- |
|  | Total viable cells | Spore titer | Total viable cells | Spore titer | Total viable cells | Spore titer | Total viable cells | Spore titer |
| WT | $1.6 \times 10^8 \pm 7.4 \times 10^7$ | $7.5 \times 10^3 \pm 9.8 \times 10^2$ | $1.7 \times 10^8 \pm 8.7 \times 10^7$ | $1.0 \times 10^4 \pm 1.2 \times 10^3$ | $8.5 \times 10^7 \pm 1.1 \times 10^7$ | $1.3 \times 10^5 \pm 5.1 \times 10^4$ | $1.0 \times 10^7 \pm 1.3 \times 10^6$ | $1.7 \times 10^5 \pm 5.8 \times 10^4$ |
| $\Delta csiA$ | $1.6 \times 10^8 \pm 2.8 \times 10^7$ | $3.1 \times 10^4 \pm 5.3 \times 10^3$ | $1.3 \times 10^8 \pm 6.5 \times 10^7$ | $8.5 \times 10^4 \pm 6.2 \times 10^4$ | $3.0 \times 10^7 \pm 2.1 \times 10^7$ | $1.5 \times 10^6 \pm 1.0 \times 10^5$ | $3.0 \times 10^7 \pm 9.8 \times 10^6$ | $2.2 \times 10^6 \pm 1.2 \times 10^6$ |
| $csiA^C$ | $1.6 \times 10^8 \pm 7.1 \times 10^6$ | $1.8 \times 10^4 \pm 7.8 \times 10^3$ | $2.3 \times 10^8 \pm 1.4 \times 10^8$ | $1.5 \times 10^4 \pm 6.8 \times 10^3$ | $3.0 \times 10^7 \pm 1.7 \times 10^7$ | $7.8 \times 10^4 \pm 2.0 \times 10^4$ | $1.0 \times 10^7 \pm 5.3 \times 10^6$ | $6.0 \times 10^4 \pm 2.0 \times 10^4$ |

We then examined cultures of the *csiA* mutant after 14h of incubation on sporulation liquid medium (SM). SM has been shown to efficiently induce sporulation in strain 630Δ*erm* and is well suited for kinetics analyses and fluorescence microscopy (Pereira *et al*., 2013, Wilson *et al*., 1982). The cells were stained with the lipophilic membrane dye FM4-64, which allows identification of the stages of sporulation up to engulfment completion, prior to imaging by phase contrast and fluorescence microscopy (Pereira *et al*., 2013). As shown in Fig. S3A, the *csiA* mutant presents more cells with signs of sporulation than the wild-type strain as determined by fluorescence and phase-contrast microscopy (630 Δ *erm*,29%; Δ *csiA*, 69%).

To rule out that the increased sporulation phenotype came from differences in vegetative growth, the wild-type and the *csiA* mutant strains were grown in SM and the optical density of the cultures was measured at 600 nm (OD_600_) at two-hour intervals for 20 hours after inoculation. Both strains showed similar growth patterns and, after about 10 hours, both entered in stationary phase (Fig. S3B). Therefore, the *csiA* mutation does not affect growth of *C. difficile* under the conditions tested.

To determine whether the kinetics of sporulation was altered in the *csiA* mutant, we analysed the formation of heat resistant spores along time. Strains were grown in SM broth, and at 12, 24, 48 and 72 hours the titer of heat resistant spores/ml of culture was determined as described in the materials and methods section (Fig. 2A and Table 1). In line with earlier results (Pereira *et al*., 2013), the titer of spores for the wild-type strain increased from 7.5×10^3^ spores/ml at hour 12, to 1.0 ×10^4^ spores/ml at hour 24, 1.3 ×10^5^ spores/ml at hour 48 and 1.7×10^5^ spores/ml at hour 72 (Fig. 2A and Table 1). As expected from the previous results, the *csiA* mutation did not affect cell viability (Table 1). In contrast, the titer of heat resistant spores in the mutant strain was higher than the wild-type for all time points: 3.1×10^4^, 8.5×10^4^ spores/mL, 1.5×10^6^ and 2.2×10^6^ spores/mL at hour 12, 24, 48 and 72, respectively (Fig. 2A and Table 1). To determine whether the *csiA* deletion was responsible for the mutant phenotype, sporulation was assessed in the complemented strain (Table 1). As shown in Fig. 2A and Table 1, complementation restored the wild-type kinetics of spore formation (1.8×10^4^, 1.5×10^4^, 7.8×10^4^ and 6.0×10^4^ spores/ml at hour 12, 24, 48 and 72, respectively). Thus, the deletion of *csiA* is responsible for the increased sporulation frequency of the mutant.

**Figure 2.**
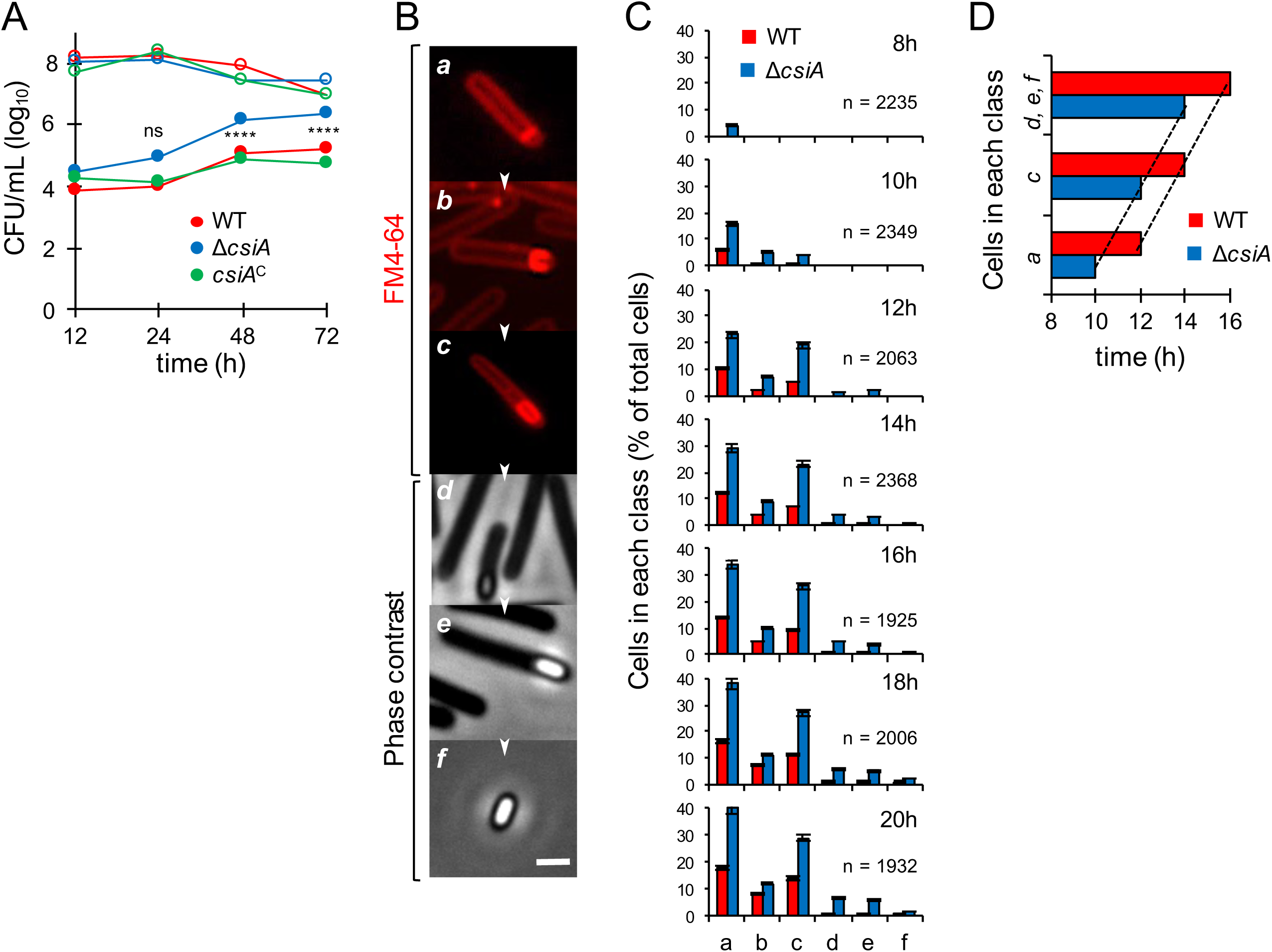
Sporulation dynamics of the Δ*csiA* mutant. **A:** Cells were grown in liquid SM and the titer of heat resistant spores (filled symbols) and total viable cells (open symbols) measured 12, 24, 48 and 72 hours following inoculation. WT, triangles; Δ*csiA* mutant, circles; *csiA^C^* (complementation strain), squares. The date in the graph represent the mean of three independent experiments (see also Table 1). Asterisks indicate statistical significance determined with a two-way ANOVA (ns, no significant; **** p<0.0001). **B:** Samples of an SM liquid culture of the WT strain were collected at 8, 10, 12, 14, 16, 18 and 20 hours after inoculation, stained with the membrane dye FM4-64 and examined by phase contrast or fluorescent microscopy. The panels illustrate the sequence of the stages in sporulation. **C:** Quantification of the percentage of cells in the morphological classes represented in B (*a* to *f*), relative to the total viable cell population, for the WT (red) and Δ *iA* mutant (blue) at the indicated times following inoculation in liquid SM. The data represent the mean ± SD of three independent experiments. The total number of cells scored (n) is indicated in each panel. **D:** Time, in hours, when at least 10% of WT or Δ*csiA* population reached the indicated stages of sporulation. The top columns shows phase-bright, phase-grey spores or free spores (classes *d*, *e* and *f*), the middle columns engulfment completion (*c*), and the bottom columns asymmetric septation (*a*).

A second frequently used method to induce sporulation is growth on 70:30 agar plates, in which the rate of sporulation is higher (Putnam *et al*., 2013). We also tested the sporulation efficiency of the *csiA* mutant in this medium. When sporulation was induced by this method, the *csiA* mutant shows a percentage of sporulation similar to the wild-type strain (Fig. S4A). Thus, the culture medium influences the *csiA* mutant phenotype (see also below).

### *csiA* negatively impacts sporulation

We next followed progress through the morphological stages of sporulation in SM, by phase contrast and fluorescence microscopy (Pereira *et al*., 2013). The wild-type and the mutant strain were grown in SM and samples were collected 8, 10, 12, 14, 16, 18 and 20 hours after inoculation. The cells were stained with the lipophilic membrane dye FM4-64 prior to microscopic examination. Cells representative of several distinctive morphological classes are shown on Fig. 2B. The results shown in Fig. 2C demonstrate that 8 hours after inoculation a significant fraction of the *csiA* cells had undergone polar division (class *a*). In contrast, class *a* cells were detected for the wild-type strain 2 hours later than in the mutant. Moreover, phase bright spores (class *e*) were observed at 14 hours for the mutant and at 18 hours for the wild-type strain.

To test whether spore morphogenesis proceeded faster in the *csiA* mutant, we determined the time at which at least 10% of the cell population had undergone polar division (class *a*), initiated engulfment (class *b*), showed complete engulfment of the forespore by the mother cell (class *c*), or presented phase-gray or phase-bright spores (classes *d* and *f*). The results in Fig. 2D show that progress through these stages of sporulation occurred at the same pace for the two strains. We conclude that the *csiA* mutant initiated sporulation earlier than the parental wild-type or that the fraction of cells entering sporulation at the end of growth is larger for the mutant. In any event, CsiA negatively affects the entry into sporulation.

### CsiA accumulates during growth independently of Spo0A

We further investigated accumulation of CsiA during growth and sporulation in SM by immunoblot analysis using an anti-CsiA antibody (Fig. 3A). CsiA starts to accumulate early during growth and is present also during entry into stationary phase when the cells start to sporulate. No signal is detected in the *csiA* mutant, while its accumulation is restored in the complementation strain, confirming the specificity of the antibody. CsiA accumulation is independent of Spo0A, the master regulator of sporulation (Fig. 3B). These results show that CsiA starts to accumulate during vegetative growth, independently of the main sporulation transcription regulators.

**Figure 3.**
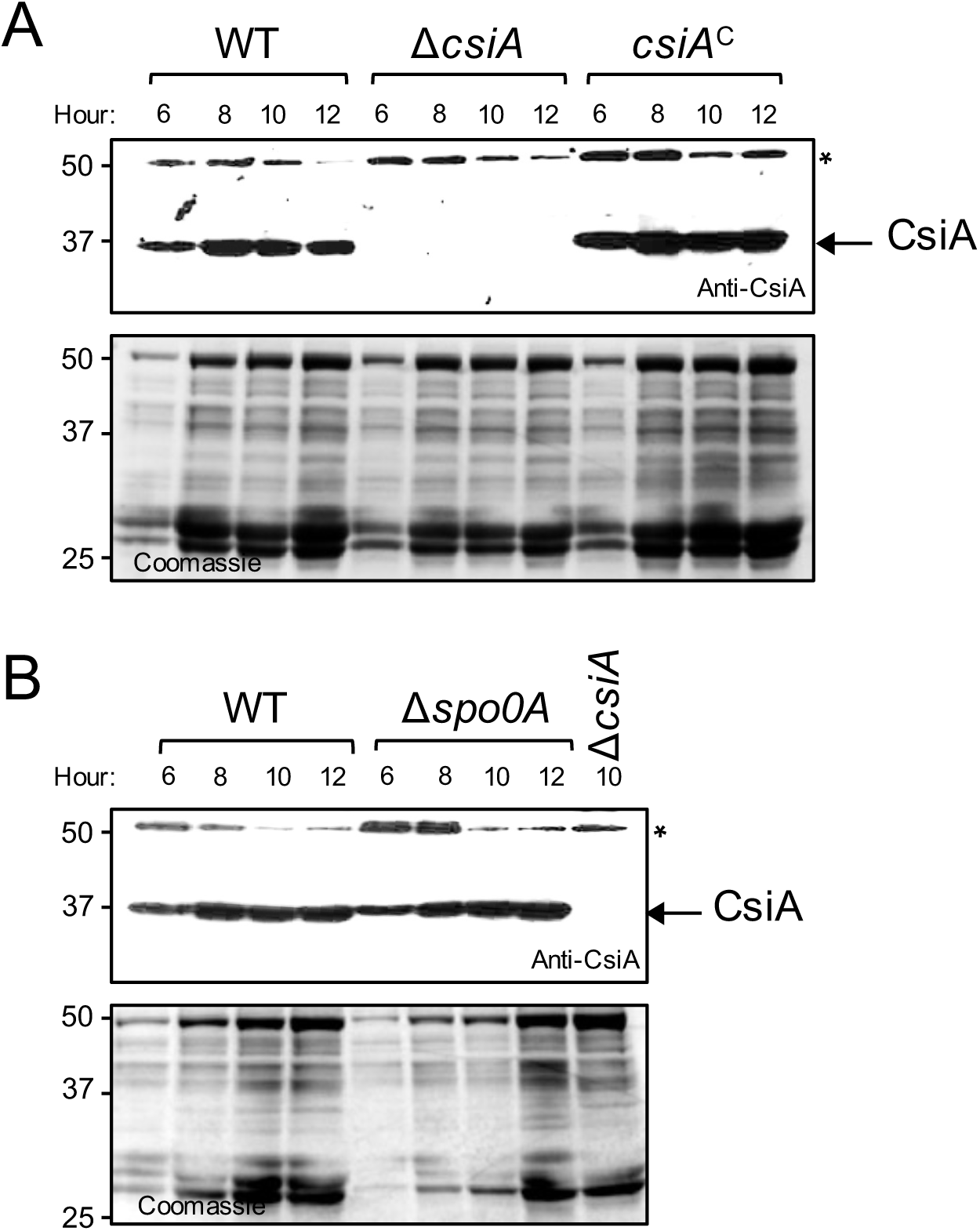
Accumulation of CsiA during growth and sporulation. The wild type strain (WT), the Δ*csiA* mutant, the complementation strain (*csiA*) (**A**) and the *spo0A* mutant (**B**) were grown in SM and samples were collected at 6, 8, 10 and 12 hours after inoculation for Western blot analysis using anti-CsiA (* cross-reactive species). The position of molecular weight markers (in kDa) is indicated on the left side of the panels.

### Deletion of *csiA* increases transcription of *spo0A*

Since Spo0A is the master regulatory protein governing entry into sporulation (Pettit *et al*., 2014, Rosenbusch *et al*., 2012, Underwood *et al*., 2009), we wanted to test whether deletion of *csiA* increased the expression of the *spo0A* gene in individual cells. We constructed a fusion of the *spo0A* promotor region to the *SNAP* reporter and we introduced the P*_spo0A_-SNAP* fusion in the *csiA* mutant and in the parental wild-type strain (Table S1). To monitor production of SNAP, samples of cultures expressing the transcriptional fusion were collected 10 hours after inoculation in SM and the cells were labelled with the TMR-Star SNAP substrate (Fig. 4A). Expression of P*_spo0A_-SNAP* was detected in 100% of the cells for both the wild-type and the *csiA* mutant (Fig. 4A). Quantification of the fluorescence signal per cell (in arbitrary units, or AU; Fig. 4B), however, shows that transcription from the *spo0A* promoter occurred at a lower intensity in the wild-type (average signal, 137.6 AU) as compared to the *csiA* mutant (average signal, 277.3 AU) (Fig. 4B).

**Figure 4.**
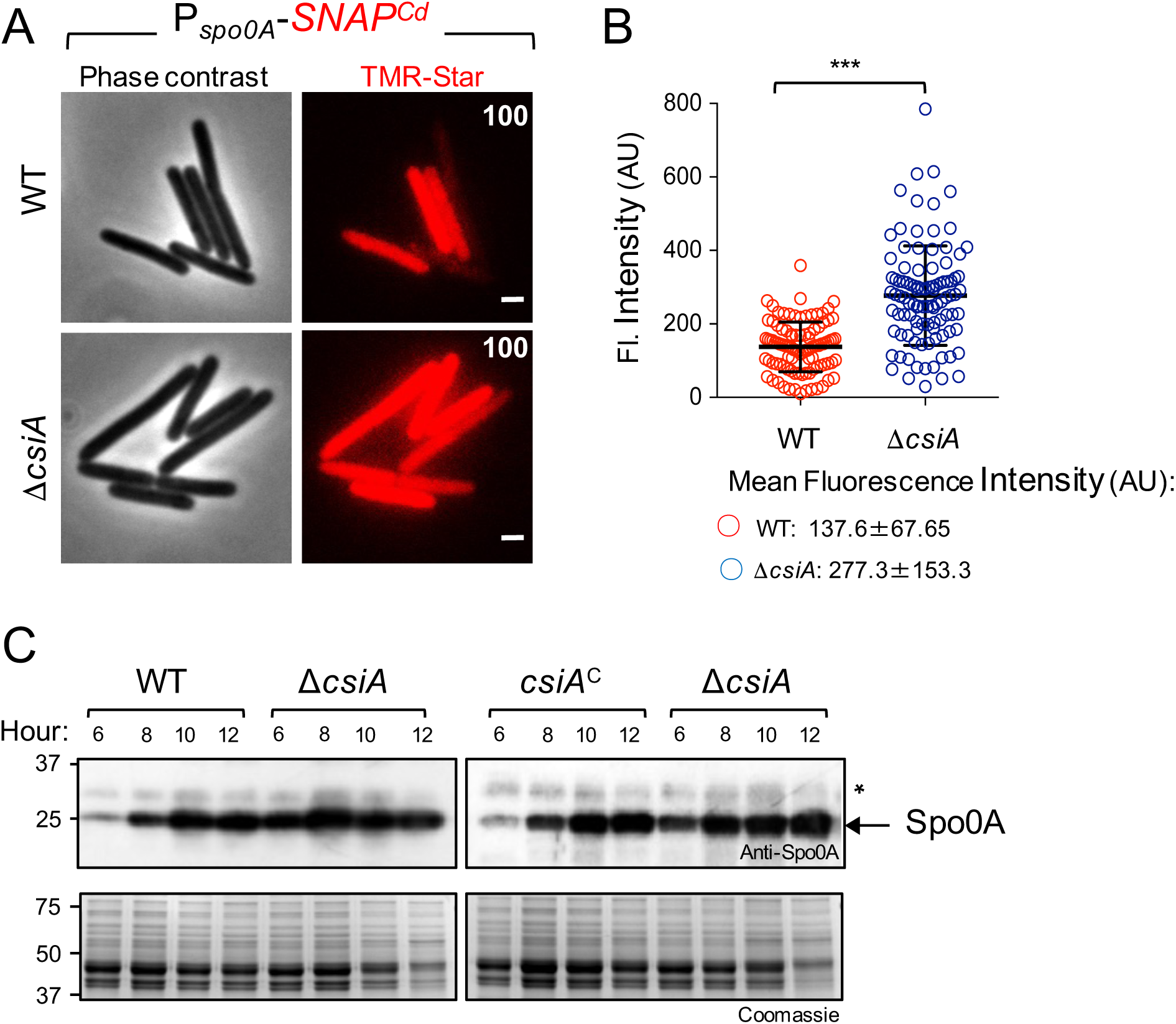
Increased expression of *spo0A* in the Δ*csiA* mutant. **A:** Microscopy analysis of *C. difficile* cells carrying a P*_spoOA_*-*SNAP^Cd^* transcriptional fusion in the WT and congenic Δ*csiA* mutant. The cells were collected after 10 h of growth in SM, stained with TMR-Star, and examined by fluorescence microscopy to monitor SNAP production. The panels are representative of the expression patterns observed. The numbers refer to the percentage of cells showing SNAP fluorescence. Data shown are from one experiment, representative of at least three independent experiments. Scale bar, 1 µm. **B:** Quantitative analysis of the fluorescence intensity (Fl.) for single cells with no signs of sporulation (n = 100 cells) of the two strains in A. Data shown are from one experiment, representative of at least three independent experiments. The numbers in the legend represent the mean and the SD of fluorescence intensity. ***, p<0.001. **C:** Samples were collected from the wild type strain (WT), the Δ complementation strain (*csiA*^C^) grown in liquid SM, at the indicated times after inoculation. Extracts were prepared and proteins (15 µg) resolved by SDS-PAGE and subjected to immunobloting using an anti-Spo0A antibody. The position of molecular weight markers (in kDa) is indicated on the left side of the panels.

To investigate whether the increased transcription from the *spo0A* promoter could also lead to increased accumulation of Spo0A, we compared the levels of Spo0A throughout sporulation by immunoblot analysis using a previously described anti-Spo0A antibody ((Serra *et al*., 2014); Fig. 4C). We found that Spo0A is detected as early as 6 hours following inoculation in SM for all strains, but that in the *csiA* mutant Spo0A accumulates at greater levels than in the wild-type or the complementation strain (Fig. 4C). Together, these results suggest that more cells of the mutant reach a threshold level of Spo0A that triggers sporulation, and that CsiA antagonizes expression of *spo0A* and/or the activity of Spo0A, since *spo0A* is subject to positive auto-regulation (Rosenbusch *et al*., 2012).

Since we did not observe a phenotype for the *csiA* mutant when sporulation was induced on 70:30 medium (see above), we compared the expression of *spo0A* in SM with that in 70:30 medium using fluorescence microscopy and single cell analysis (Fig. S3B). Quantification of the fluorescence signal per cell (Fig. S4B) shows that transcription from the *spo0A* promoter is higher in 70:30 medium in both the wild-type (average signal, 267.1 AU) and *csiA* mutant (average signal, 332.8 AU), reaching levels similar to those observed in the mutant when sporulation is induced in SM (average signal,372 AU) (Fig. S4B). This result suggests that in 70:30 medium, the wild-type strain reaches the threshold levels of *spo0A* required for sporulation in a higher number of cells than in SM, similar to what was observed for the *csiA* mutant in both media. Hence, CsiA acts as a repressor only under certain culturing conditions that induce sporulation, with the effect of the *csiA* mutation being masked in 70:30. medium

### Phosphorylated Spo0A accumulates at higher levels in the *csiA* mutant

Spo0A is activated by phosphorylation (Fujita & Sadaie, 1998, Lewis *et al*., 1999). The activated Spo0A protein, Spo0A∼P, binds to DNA promoter regions containing a Spo0A-binding motif and then regulates the expression of Spo0A-dependent genes. Since spo0A is auto-regulatory, and higher expression per cell is seen in the *csiA* mutant in SM, we asked whether Spo0A∼P would accumulate at higher levels in the mutant. For this, we used Phos-tag SDS-PAGE, where migration of Spo0A∼P is delayed, and the two forms (phosphorylated and unphosphorylated) are then detected by western blot using anti-Spo0A antibodies. The wild-type and the mutant strain were grown in SM and samples were collected 6, 8, 10 and 12 hours after inoculation. Whole cell extracts were prepared and the proteins resolved by Phos-tag SDS-PAGE (Fig. 5A). Two bands were detected in this manner. The upper band disappeared when the samples were boiled prior to electrophoretic resolution, which indicates that the upper band corresponds to Spo0A∼P and the lower band to the unphosphorylated form (Yasugi *et al*., 2016).

**Figure 5.**
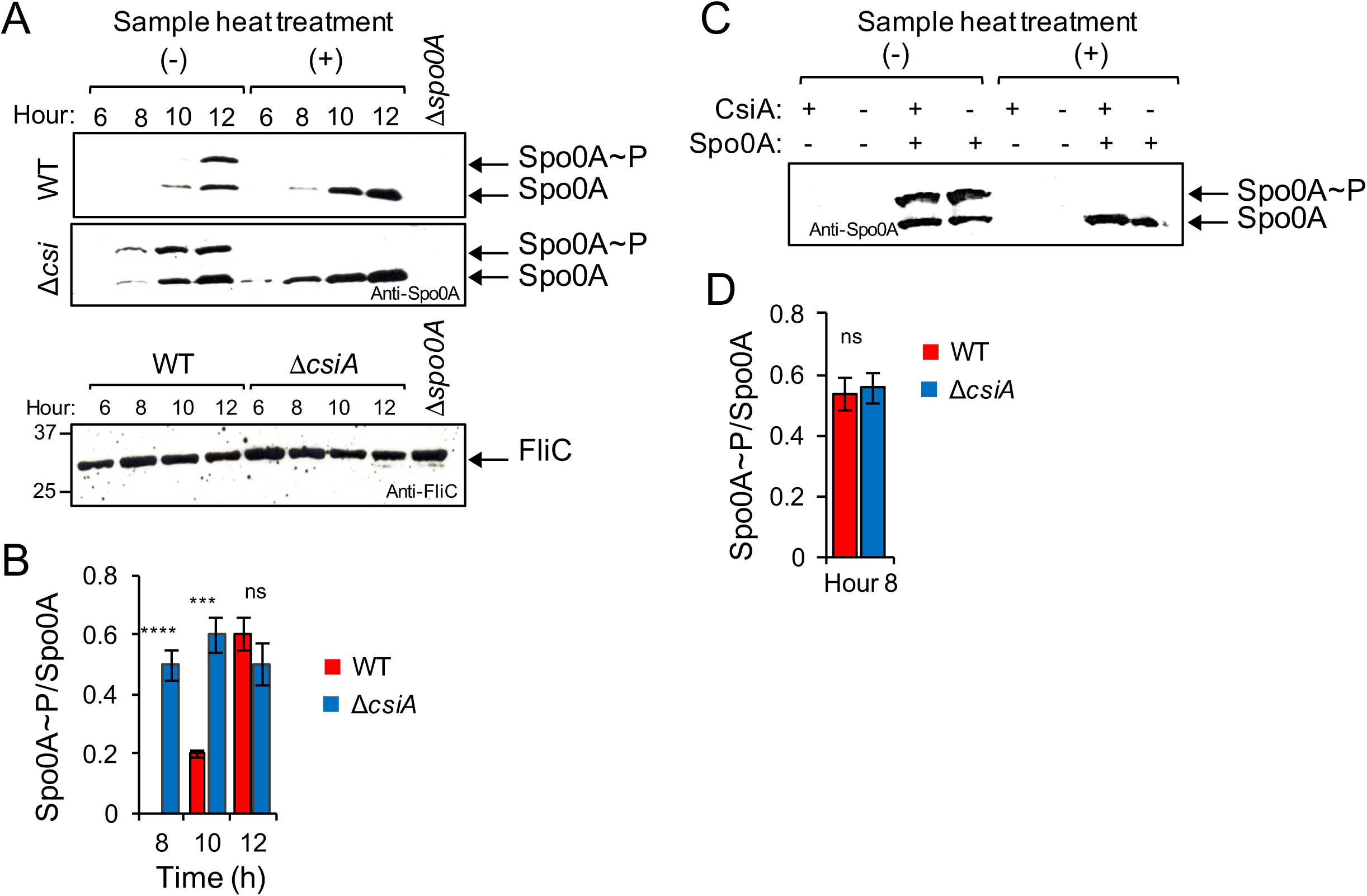
Phosphorylation of Spo0A in the Δ*csiA* mutant. **A:** Samples were collected from the WT and the Δ*csiA* mutant grown in liquid SM, at the indicated times after inoculation. Extracts were prepared and proteins (15 µg) resolved by Phos-tag SDS-PAGE and subjected to immunobloting using an anti-Spo0A antibody. In the bottom panel the same extracts were loaded in a SDS-PAGE and subjected to immunoblotting using an anti-FliC antibody, as a loading control. The position of molecular weight markers (in kDa) is indicated on the left side of the panels. **B and D:** The ratio of Spo0A∼P to Spo0A was assessed with Image J software and represented for the WT and the Δ*csiA* mutant at the indicated times (in hours) after inoculation in liquid SM. All data represent the means ± SD from three independent experiments. Asterisks indicate statistical significance determined by two-tailed Student t-test (ns, no significant; **** p<0.0001, *** p < 0.001). **C:** Extracts were prepared from liquid SM cultures of the P*_tet_*-*spo0A* and Δ*csiA* P*_tet_*-*spo0A* strains grown in the presence (50 nM) or in the absence of anhydrotetracycline, 8 hours after inoculation. Proteins (15 µg) were resolved by Phos-tag SDS-PAGE and subject to immunoblotting using an anti-Spo0A antibody. In **A** and **C**: The faster migrating bands (black arrows) show the unphosphorylated form of Spo0A (Spo0A), and the slower migrating bands indicate the phosphorylated form of Spo0A (Spo0A∼P; red arrows). The samples heated at 100°C for 5 min were loaded as a control for the position of unphosphorylated Spo0A.

We found that in the *csiA* mutant, both forms of Spo0A accumulate earlier and at higher levels than in the wild-type. We then quantified band intensities in order to estimate the ratio of the phosphorylated to unphosphorylated form of Spo0A (Spo0A~P/Spo0A) using Image-J software. The Spo0A~P/Spo0A ratio was higher at hour 8 and 10 in the mutant as compared to the wild-type strain (Fig. 5B).However, at hour 12 the ratio between the two forms was similar between the two strains.

This result suggests that the level of Spo0A phosphorylation is directly proportional to the level of the protein. Together with the observation that *spo0A* is expressed in most if not all of the pre-divisional cells (Fig. 4A), our findings suggest that phosphorylation of Spo0A is not a limiting factor for the initiation of sporulation. Possibly then, the level of *spo0A* per cell could limit entry into sporulation.

### CsiA acts at the level of *spo0A* transcription

Since we found increased transcription of P*_spo0A_-SNAP^Cd^* in the *csiA* mutant, we reasoned that the increased sporulation in the mutant could be due to increased transcription or to increased activity of Spo0A. To uncouple transcription from activation of Spo0A by phosphorylation, we placed the *spo0A* gene under the control of the anhydrotetracycline (ATc) responsive P*_tet_* promoter (Dembek *et al*., 2017). In this way Spo0A could accumulate to similar levels in the wild-type and in the *csiA* mutant strains, allowing us to test whether increased levels of Spo0A∼P accumulated in the mutant. The strains were grown in SM supplemented with 50 nM of ATc, a concentration at which Spo0A accumulates to levels similar to the wild-type strain (Dembek *et al*., 2017), and proteins in whole cells extracts were resolved by Phos-tag SDS-PAGE (Fig. 5C). In both strains Spo0A accumulates at similar levels and the Spo0A∼P/Spo0A ratio was similar (Fig. 5C and D). In agreement with this observation, heat test revealed no difference in the number of heat resistant spores produced by both strains (Table 2). Together these results suggest that CsiA acts mainly by decreasing the levels of *spo0A* transcription per cell.

**Table 2.** Sporulation efficiency of the Ptet-spo0A alleles. The total viable cell count and the spore titer was determined 24 hours following inoculation into sporulation medium supplemented with 50 nM of anhydrotetracycline. The heat resistant spore count by determining the cfu/mL obtained after treatment at 70°C. The results shown are averages and standard deviations for three biological replicates.

|  | 24h |  |
| --- | --- | --- |
|  | Total viable cells | Spore titer |
| WT *P<sub>tet</sub>-spo0A* | $2.1 \times 10^7 \pm 1.3 \times 10^6$ | $6.8 \times 10^6 \pm 1.3 \times 10^5$ |
| Δ*csiA* *P<sub>tet</sub>-spo0A* | $2.2 \times 10^7 \pm 1.6 \times 10^6$ | $7.6 \times 10^6 \pm 1.7 \times 10^5$ |

### The *csiA* mutant shows increased expression of Spo0A-dependent genes

The increased expression of *spo0A* and the increased sporulation of the *csiA* mutant suggested that a larger fraction of the population entered the sporulation pathway. To independently test this idea, we analysed the expression of the genes coding for the early sporulation sigma factors *sigE* and *sigF*, whose transcription is under Spo0A control. To monitor transcription of *sigE* and *sigF*, we used transcriptional fusions of the *spoIIA* (driving σ^F^ production) and *spoIIG* (driving σ^E^ production) promoters to the SNAP^Cd^ reporter (Pereira *et al*., 2013). Samples of cultures expressing each of the promoter-SNAP^Cd^ fusions were collected 14 hours after inoculation in SM and the cells labelled with TMR-Star and the membrane dye MTG, to allow identification of the different stages of sporulation (Fig. 6A). Expression of both *sigF* and *sigE* was first detected in pre-divisional cells and cells that had undergone polar division, as reported previously (Pereira *et al*., 2013).

**Figure 6.**
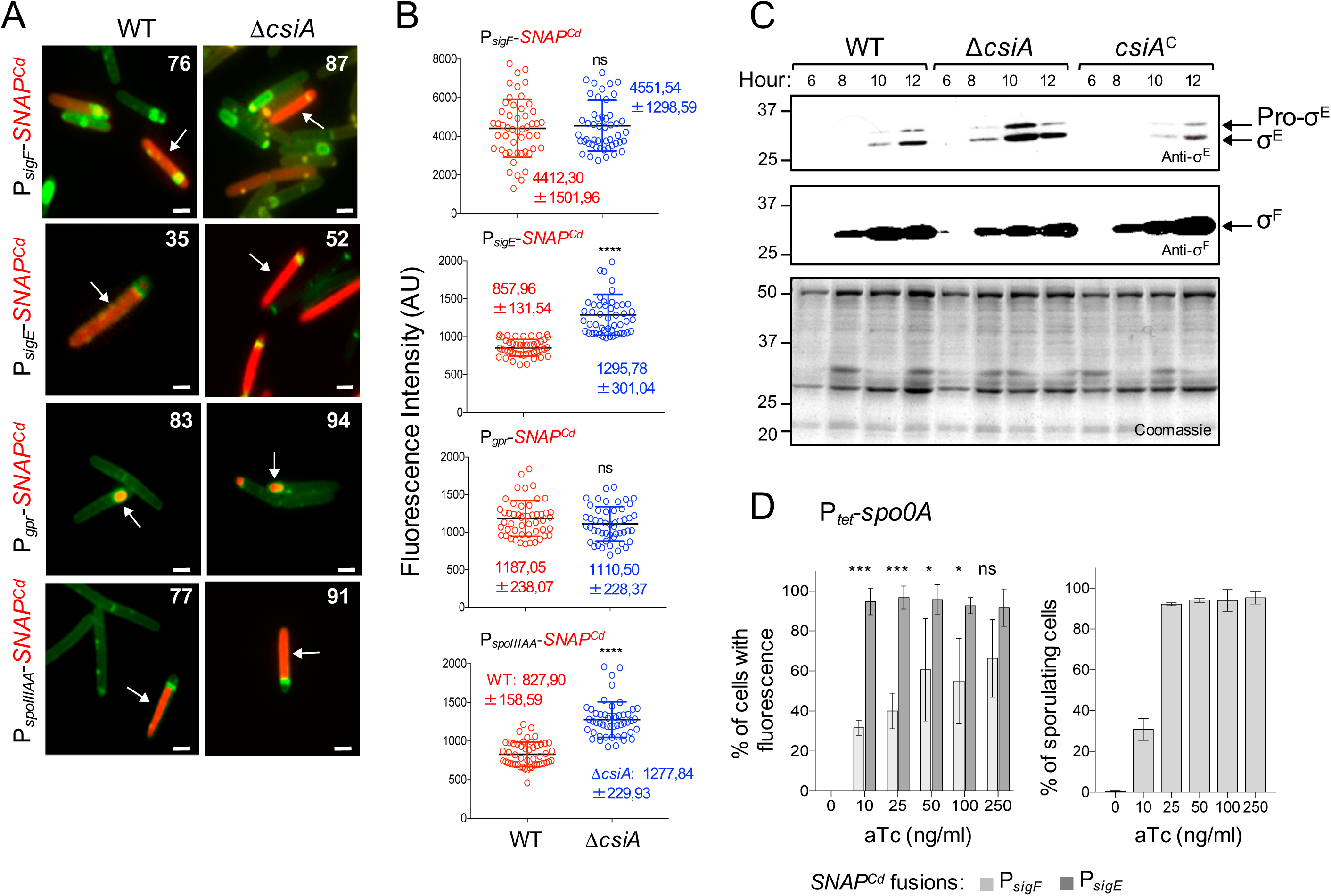
Increased sporulation-specific gene expression in the Δ*csiA* mutant. **A:** Microscopy analysis of *C. difficile* cells carrying transcriptional fusions of the *sigF*, *sigE, gpr* and *spoIIIA* promotors to SNAP in the WT and in the Δ*csiA* mutant. The cells were collected after 12 h of growth in liquid SM, stained with TMR-Star and the membrane dye MTG, and examined by fluorescence microscopy to monitor SNAP production. The merged image shows the overlap between the TMR-Star (red) and MTG (green) channels. The images are representative of the expression patterns observed. The numbers refer to the percentage of cells showing SNAP fluorescence. Data shown are from one experiment, representative of at least three independent experiments. Scale bar, 1 µm. **B:** Quantitative analysis of the fluorescence intensity (in arbitrary units, AU) of the reporter strains for *sigF*, *sigE, gpr* and *spoIIIA* transcription, as indicated. The numbers in the legend represent the average and the standard deviation of fluorescence intensity for the cells considered (n = 50 cells). ***, p<0.0001. **C:** Extracts were prepared from liquid SM cultures of the WT, Δ*csiA* mutant and the complementation strain (*csiA^C^)*, grown, at the indicated times after inoculation. Proteins (15 µg) were resolved by SDS-PAGE (12%) and subject to immunoblotting with anti-σ and anti-σ antibodies. The position of molecular weight markers (in kDa) is indicated on the left side of the panels. **D:** measurement of the increase in Spo0A activity as a function of aTc using a P*_sigF_*-*SNAP* (light grey) or P*_sigE_*-*SNAP* (dark grey) reporter and the formation of asymmetric septa. Asterisks indicate statistical significance determined by two-way ANOVA (ns, no significant, *** p < 0.001, * p<0.01).

Quantification of the fluorescence signal shows that although for *sigF* the percentage of cells with a fluorescence signal is higher in the *csiA* mutant (52%) than in the wild-type strain (35%), the average intensity of the signal did not differ much between the two strains (1187.05 AU for the wild type and 1110.50 AU for the mutant) (Fig. 6B). Transcription of *sigE*, however, was lower in the wild-type, with 76% of the cells showing a signal with an average intensity of 857.96 AU, as compared to the mutant, in which 87% of cells showed a fluorescence signal with a significantly higher average intensity of 1295.78 AU (Fig. 6B).

We next examined the effect of the *csiA* mutant on the activity of σ^F^ and σ^E^ by monitoring expression of transcriptional fusions of the *gpr* (controlled by σ^F^) and *spoIIIA* (controlled by σ^E^) promoters to the SNAP^Cd^ reporter (Pereira *et al*., 2013). Expression of *gpr* was detected in 83% of the sporulating cells of the wild-type and in 94% of the *csiA* sporangia (Fig. 6A), while expression of *spoIIIA* was detected in 77% of the sporulating cells in the wild-type and in 91% of the *csiA* sporangia (Fig. 6A). A quantitative analysis revealed no effect of the *csiA* mutation on *gpr* expression (average intensity of 4412.30 AU for the wild type and 4551.54 AU for the mutant; Fig. 6B). In contrast, the average intensity of the fluorescence signal from P*_spoIIIA_*-*SNAP^Cd^* was higher in cells of the mutant (1277.84 AU for the mutant) as compared to the wid-type (827.90 AU) (Fig. 6B). This increase likely results from the augmented expression of *sigE* in the *csiA* mutant (see above).

### Earlier accumulation and processing of pro-**σ**^E^ in the *csiA* mutant

The increased activity of σ^E^, but not of σ^F^, suggested to us that σ^E^ could accumulate to higher levels in cells of the *csiA* mutant as compared to the wild-type and that in contrast, the levels of σ^F^ would be similar in the mutant and the wild-type. To test this prediction, we monitored the accumulation of σ^F^ and σ^E^ by immunoblotting during entry into sporulation. The mature, active form of σ^E^ was first detected 8 hours after inoculation in the mutant and only at 10 hours in the wild-type (Fig. 6C). Importantly, we found that the accumulation of the mature form of σ^E^ is also higher for the *csiA* mutant as compared to the wild-type, with the protein accumulating at levels 8 times higher in the mutant 10 hours after inoculation (Fig. 6C). On the other hand, there was no significant difference on the accumulation of σ^F^ between the wild-type and the mutant. Thus, the lack of CsiA leads to premature accumulation and activation of pro-σ^E^, but does not affect the accumulation of σ^F^.

### Expression of *sigE* establishes a threshold for sporulation

Higher levels of Spo0A∼P accumulate in the *csiA* mutant, leading to increased expression of the Spo0A-dependent genes. Consequently, the higher levels of Spo0A∼P increase the number of cells showing asymmetric division (Fig. 2C), and on transcription from the *spoIIG* promoter, but not from the *spoIIA* promoter (Fig. 6B). These results strongly suggest that asymmetric division and the level of active σ^E^ are a limiting factor for sporulation. To test this idea we inserted the P*_sigF_*- and P*_sigE_*-*SNAP^Cd^* fusions in the strain in which *spo0A* is under the control of P*_tet_* promoter (Dembek *et al*., 2017). The strains were grown in SM to an OD_600_ of approximately 0.2, supplemented with a range of ATc concentrations, from 0– 250 ng/ml, and 2 hours after addition the cells were labelled with TMR. In the absence of the inducer, no fluorescence was detected in either strain. In contrast, a clear dose-dependent response in the percentage of cells showing fluorescence from the P*_sigE_*-*SNAP* fusion was observed at concentrations of inducer from 0 to 50 ng/ml (Fig. 6D), reaching a plateau at higher aTc concentrations (approximately 70% of the cells). Interestingly, this is also the concentration at which wild type levels of heat resistance spores are reached in the strain carrying *spo0A* under P*_tet_* control (Dembek *et al*., 2017). In contrast, low levels of *spo0A* (10 ng/ml) are required to detect expression of *sigF* in nearly all of the cells in the population (approximately 95%; Fig. 6D). Since the formation the polar septum is also governed by Spo0A∼P (Ben-Yehuda & Losick, 2002, Levin & Losick, 1996), we also measured the number of cells showing asymmetric septa in the P*_tet_*-*spo0A* strain at the same range of ATc concentrations, following staining of the cells with the membrane dye FM4-64 prior to microscopic examination (Fig. 6D). At 10 ng/ml of aTc only approximately 35% of the cells showed asymmetric septa or were at intermediates stages during engulfment; this percentage increased to more than 90% at 25 ng/ml of aTc.

Thus, our results suggest that there are different Spo0A∼P thresholds for the formation of the asymmetric septum, and for the activation of *spoIIA* and *spoIIG* transcription. At least under some conditions, a fraction of the cells that activate *spoIIA* expression may not reach a point that allows *spoIIG* expression or asymmetric division.

### Deletion of *csiA* mainly affects sporulation

Spo0A regulates directly or indirectly a number of virulence-associated factors such as motility, biofilm formation and toxin production (Pettit *et al*., 2014, Rosenbusch *et al*., 2012, Underwood *et al*., 2009). Therefore, we wanted to determine whether the *csiA* mutation had a more global effect on gene expression, for that we decided to compare the transcriptome of the mutant with the wild type after 10 hours of growth in SM. We used two biological replicates and genes were considered differentially expressed if the fold change was > 2 and the adjusted p value <0.01. We found that 165 genes were overexpressed in the mutant, while only 21 genes were under-expressed (Table 3). From these genes, 69% are genes regulated by Spo0A or by sporulation-specific cell type-specific sigma factors (Fig. 7A and Table 3). No toxin, biofilm or motility genes are among these genes (see also Fig. S5).

**Figure 7.**
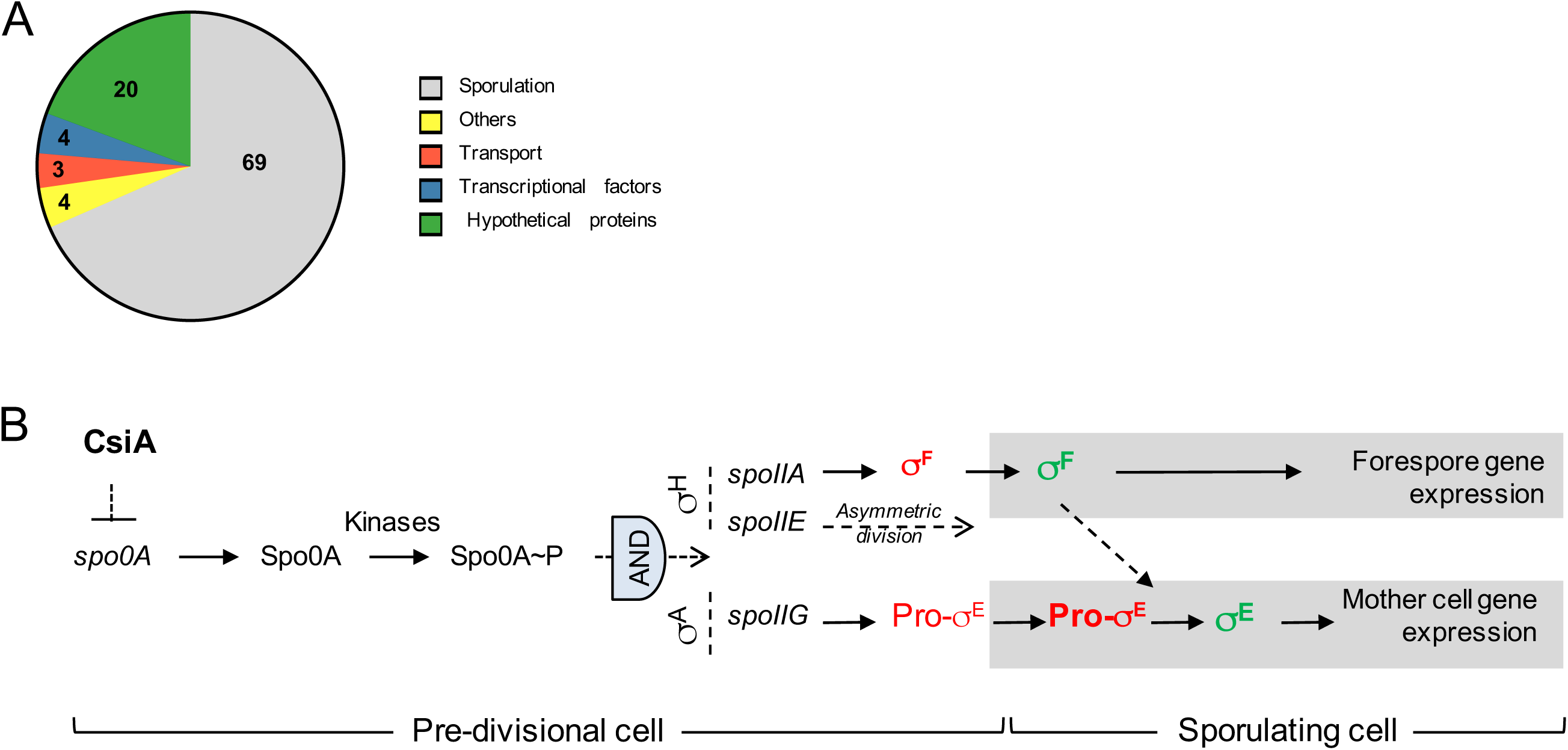
Model for the function of the *csiA* gene. **A:** Functional classes of the genes that are up-regulated in the Δ*csiA* mutant based on RNA seq data (left; the number represent the percentage of each functional class) and the up-regulated sporulation genes in the indicated cell type-specific regulons (right; the numbers represent the percentage of each of the regulons). See also Table 3 **B.** Model for the function of the *csiA* gene in controlling sporulation initiation in *C. difficile*.

## Discussion

A genomic signature of sporulation in the human intestinal microbiome (Browne *et al*., 2016) includes not only genes with an established function in sporulation (Fig. 1) but also a group of 20 genes of unknown function. We hypothesized that these uncharacterized genes are important for sporulation. To test this hypothesis, we characterized a mutant in the *csiA* (*CD25890*) gene. We show that the CsiA protein accumulates during growth in a medium in which sporulation is induced, and also during sporulation (Fig. 3). Attempts to localise *csiA* expression during sporulation using a SNAP transcriptional fusion to the 331 bp region upstream of *csiA* failed, even if this same region is sufficient to allow the expression of *csiA* at the *pyrE* locus at levels that permit complementation of the *csiA* deletion mutation.

We found that under certain culturing conditions, *csiA* deletion has an impact on sporulation. The *csiA* mutant reaches a higher titer of spores than the wild type strain when grown in sporulation medium (SM). We show here that in the *csiA* mutant spore differentiation is triggered in a larger fraction of the population (Fig. 2). This mutant maintains high levels of *spo0A* expression and shows higher expression of genes required for sporulation, such as *spoIIG* (Fig. 4 and 6). Since Spo0A is auto-regulatory, the transcriptional fusion reports both transcription of *spo0A* as well as the activity of Spo0A which requires phosphorylation of the protein (Rosenbusch *et al*., 2012). We show that higher levels of Spo0A∼P accumulate in the *csiA* mutant earlier during growth, leading to increased expression of Spo0A-dependent genes. Uncoupling of transcription from activation by using an inducible promoter to control *spo0A* expression showed that CsiA does not act on the phosphorylation of Spo0A but rather at the level of *spo0A* transcription (Fig. 5). In cells growing in SM, Spo0A∼P is directly proportional to the total level of protein, indicating that under these conditions phosphorylation is not a limiting step for the initiation of sporulation.

Our finding that higher accumulation of Spo0A correlates with enhanced sporulation contrasts with studies performed in *B. subtilis*, where accelerated accumulation of Spo0A independent of the phosphorelay was detrimental to sporulation (Fujita & Losick, 2005, Vishnoi *et al*., 2013). Rather, Spo0A needs to accumulate in a gradual manner to trigger sporulation and this requires its phosphorylation through the phosphorelay (Vishnoi *et al*., 2013). This suggests that a gradual build-up of Spo0A may not be important for the sporulation process in *C. difficile*. Previous studies have already shown that *spo0A* expression can be increased form an inducible promoter in a dose-dependent manner, and that the level of expression correlates with sporulation efficiency, which reaches values higher than those observed in a congenic wild-type strain (Dembek *et al*., 2017).

Intriguingly, in the *csiA* mutant, higher Spo0A∼P levels impact the formation of the asymmetric septum, possibly through *spoIIE,* which is also under the control of Spo0A and is required for proper septum formation (Ben-Yehuda & Losick, 2002, Levin & Losick, 1996). The mutation also increased *spoIIG* transcription, while not affecting *spoIIA* transcription (Fig. 2 and 6). These events are under the control of Spo0A (Ben-Yehuda & Losick, 2002, Levin & Losick, 1996, Pettit *et al*., 2014, Rosenbusch *et al*., 2012, Saujet *et al*., 2011). The *spoIIA* operon (*spoIIAA*-*spoIIAB*-*sigF*) is transcribed under the joint control of the σ^H^ sigma factor and Spo0A, which binds to Spo0A boxes in the *spoIIA* promoter region (Pettit *et al*., 2014, Rosenbusch *et al*., 2012, Saujet *et al*., 2011). Our results, however, suggests that a factor or factors other than Spo0A contributes to expression of *sigF*, as also shown in *C. acetobutylicum* (Alsaker *et al*., 2004). *spoIIE* is involved in the Spo0A-dependent switch in the septum localization from mid-cell to the pole, and its expression is also σ^H^- and Spo0A-dependent (Ben-Yehuda & Losick, 2002, Khvorova *et al*., 1998, Levin & Losick, 1996, Saujet *et al*., 2011). In contrast, the *spoIIG* operon (*spoIIGA*-*sigE*) is under the control of σ^A^ and is also positively regulated by Spo0A. Therefore, differences on the role of CsiA in the expression from different promoters are unlikely to be related to the regulation of the σ factor. Rather, CsiA may affect the complexes made by σ^H^ and Spo0A and by σ^A^ and Spo0A at different promoters.

In line with the *SNAP^Cd^-*based single cell transcriptional analysis, we also found that accumulation of the active, processed form of σ^E^ is higher in the *csiA* mutant as compared to the wild-type. In contrast, there was no significant differences on the accumulation of σ^F^. This is in agreement with previous results showing that the expression of the σ regulon in the mother cell was not strictly under the control of σ (Saujet *et al*., 2013).

Previous studies have shown that the DNA-binding domain of Spo0A (Spo0A-DBD) is able to bind to the promoter regions of *sigF*, *spoIIE* and *sigE*, (Rosenbusch *et al*., 2012). Spo0A-DBD binds at low concentration to *spoIIA* and *spoIIE*, and a higher concentration is needed to detect binding to the *sigE* promoter. Here, we used a strain in which *spo0A* expression is under the control of an inducible promoter (Dembek *et al*., 2017) to test for the requirement of different amounts of Spo0A to trigger *sigF* and *sigE* expression, as well as formation of asymmetric septa (an event that is also under Spo0A control). Under the conditions tested, phosphorylation was not a limiting step for sporulation (Fig. 5). Therefore, we infer that the amount of *spo0A* expression is proportional to the level of Spo0A∼P. We found that expression from the *spoIIA* promoter increases dramatically at low levels of *spo0A* expression (10 ng/ml of the inducer aTc), while higher levels of *spo0A* expression (250 ng/ml of aTc) are required for *sigE* to be expressed in the majority of the cells. Asymmetric division is achieved in almost 100% of the cells when *spo0A* was induced with 25 ng/ml of aTc (Fig. 6). We were not able to detect *spoIIE* expression using a *SNAP* transcriptional fusion. These results indicate that as in *B. subtilis* many of the *spo0A*-dependent genes respond to the transcriptional factor in a dose-dependent manner (Fujita *et al*., 2005). In *B. subtilis* the threshold of Spo0A activity required for σ^F^ activation is below that for the activation of σ^F^ (Narula *et al*., 2012). In *B. subtilis*, cells that form an asymmetrical septum but do not activate σ^F^ or σ^F^ are able to resume vegetative growth (Dworkin & Losick, 2005); commitment to sporulation occurs only after the activation of both of the first cell type-specific σ^F^ factors, σ^F^ and σ^F^. Since spore formation is an energy consuming process and irreversible after specific points, in *C. difficile* commitment to sporulation may be postponed until σ^E^ accumulates to sufficient levels. This may be achieved by controlling *spo0A* expression, and *csiA* may be involved in this process (Fig. 7D).

The impact of *csiA* on sporulation and on expression of sporulation-specific genes is significantly influenced by the culture medium. The two sporulation conditions used here differ in medium composition, and this may indicate that the signalling to enter sporulation may be different, dependent on the environment.

The specific external signals that activate sporulation in *C. difficile* are not known, but it is anticipated that, like in *B. subtilis*, the sporulation-initiating and -inhibiting signals for *C. difficile* act on Spo0A (Childress *et al*., 2016, Underwood *et al*., 2009).

Many factors that influence the sporulation pathway have been identified. Two global regulators, CcpA and CodY, affect the initiation of sporulation by repressing the expression of sporulation-related genes in high-nutrient conditions (Antunes *et al*., 2012, Dineen *et al*., 2007, Girinathan *et al*., 2018, Nawrocki *et al*., 2016). Both of these transcription repressors also repress the expression of the toxin encoding genes (Antunes *et al*., 2012). Another negative regulator of sporulation initiation is the histidine kinase CD1492. The *CD1492* mutant shows increased expression of *spo0A*-dependent genes, but also exhibits decreased toxin production and motility. RstA, acts as a positive regulator of sporulation, although the exact nature of this effect remains unclear (Edwards *et al*., 2016). The DNA binding domain of RstA was shown to directly bind to toxin and motility genes, repressing their expression, but this domain is not required for the positive effect exerted on sporulation (Edwards *et al*., 2019). The genome of *C. difficile* also encodes for two oligopeptide permeases (Opp and App) that in *B. subtilis* are involved in the control of sporulation initiation in a cell density-dependent manner (Koide & Hoch, 1994, Perego *et al*., 1991). Given the ecological niche of *C. difficile*, cell density may not be a signal that triggers sporulation, and in contrast to *B. subtilis*, the *opp* and *app* mutants show increased sporulation-specific gene expression, and exhibit a hyper-sporulation phenotype with no impact on toxin production or motility (Edwards *et al*., 2014). It was suggested that *opp* and *app* may be involved in sporulation as a response to nutrient availability. Like *csiA*, these mutants also show an increase on the expression of the *sinRR’* operon (Table 3), previously shown to be involved in sporulation, toxin production and motility (Girinathan *et al*., 2018). It was shown that overexpression of *sinR* increased sporulation efficiency, while overexpression of *sinR’* reduced sporulation. However, to our knowledge the overexpression of the entire operon, as occurs in the *csiA* and *opp*, *app* mutants, was not tested. With the exception of the *opp* and *app* peptide transporter mutants, all the mutants in genes involved in the control of initiation of sporulation also have an impact on toxin production and/or motility.

Since deletion of *csiA* has no impact on toxin production and motility, the *csiA* function is most likely independent from SigD, RstA, CD1492, and CD2492. In addition, *csiA* must not be impacting activity of CodY or CcpA, both of which also affect toxin production.

*B. subtilis csiA* orthologue (*yloC*) is in an operon with the *remA* gene, which was also previously identified as part of a sporulation genomic signature ((Abecasis *et al*., 2013, Traag *et al*., 2013), Fig. S1, orange gene). RemA is required for activating expression of the extracellular matrix biosynthetic operons during biofilm development by undomesticated strains of *B. subtilis* (Winkelman *et al*., 2009, Winkelman *et al*., 2013). While *remA* is not present in *C. difficile* and neither YloC nor CsiA show sequence similarity to proteins of known function, the genomic context suggested a role in the regulation of genes at the onset of the stationary growth, which may have different outcomes in the two bacteria (biofilm or sporulation) due to differences on the specific signals that lead to different responses.

Our study has identified a new factor that affects sporulation initiation, CsiA, which negative impacts sporulation by affecting the expression of *spo0A*. The medium dependence of CsiA function may reflect differences in nutrient-signalling of *spo0A* expression and this observation may be explored to identify some of those signals. In all, our analysis shows that the genomic signature of sporulation within the human intestinal microbiome can be used to identify new genes important during the sporulation process. Further studies are needed to established the function of the other signature genes. This may lead to the identification of new genetic determinants for spore formation whose products could act as important targets for the design of drugs effective against all spore-formers.

## Materials and Methods

### Growth conditions and general methods

Bacterial strains and their relevant properties are listed in Table S1. The *Escherichia coli* strain DH5α (Bethesda Research laboratories) was used for molecular cloning, while the strain HB101 (RP4) was used as the donor in *C. difficile* conjugation experiments (Hussain *et al*., 2005). Luria-Bertani medium was routinely used for growth and maintenance of *E. coli.* When appropriate, ampicillin (100 µg/ml) or chloramphenicol (15 µg/ml) was added to the culture medium. The *C. difficile* strains used in this study are isogenic derivatives of the wild-type strain 630 *erm* (Hussain *et al*., 2005) and were Δ routinely grown anaerobically (5% H_2_, 15% CO_2_, 80% N_2_) at 37°C in brain heart infusion (BHI) medium (Difco) (Wilson *et al*., 1982). Assays for toxin production were done in tryptone yeast extract (30g/L tryptone; 20g/L yeast extract) (TY). When necessary, cefoxitin (25 µg/ml) and thiamphenicol (15 µg /ml) were added to *difficile* cultures. A defined minimal media (CDMM) (Karasawa *et al*., 1995) with 1% agar was used as uracil-free medium when performing genetic selections.

Sporulation was tested on liquid Sporulation Medium (SM; for 1L: Bacto tryptone 90 g, Bacto peptone 5 g, (NH_4_)_2_SO_4_ 1 g, Tris base 1.5 g, pH 7; (Pereira *et al*., 2013)) and 70:30 agar medium (for 1L: Bacto peptone 63 g, proteose peptone 3.5 g, (NH_4_)_2_SO_4_ 0.7 g, Tris base 1.06 g, brain heart infusion extract 11.1 g, yeast extract 1.5 g, cysteine 0.3 g and agar 15 g, (Putnam *et al*., 2013)). Sporulation was regularly induced by inoculating in SM liquid. To determine the total number of cells, the cells were serially diluted and plated on BHI with 0.1 % taurocholate (Sigma-Aldrich) to ensure efficient spore germination. To determine the number of spores, the cells were heat killed by incubation for 30 min at 70°C prior to plating on BHI with 0.1 % taurocholate.

### Construction of the *csiA* mutant

A *csiA* in-frame deletion mutant was generated using allele-couple exchange (ACE) in *C. difficile* 630 *erm* Δ*pyrE* as described (Ng *et al*., 2013). The homology regions upstream and downstream of the desired junction point within *csiA* were PCR-amplified and cloned into pMTL-YN3 as follows (Ng *et al*., 2013). The upstream fragment (742 bp) was amplified using the primers *csiA*_AscI_Fwd and *csiA*_LHA_Rev and the downstream fragment (735 bp) was amplified using the primers *csiA*_RHA_Fwd and *csiA*_SbfI_Rev. The fragments were then joined by overlapping PCR, the resulting fragment cleaved with AscI and SbfI and cloned between the same sites of pMTL-YN3, yielding pAM37. This plasmid was introduced into *E. coli* HB101 (RP4) and then transferred to strain 630 *erm* Δ*pyrE* by conjugation (Heap *et al*., 2007). Following two passages on BHI agar supplemented with 25 µg/mL cefoxitin and 15 µg/mL thiamphenicol, colonies that were noticeably larger (indicative of plasmid integration) were screened by colony PCR to identify single-crossover mutants using primers flanking the upstream and downstream homology regions in conjunction with a plasmid-specific primer (P3 with P2 and P4 with P1) to amplify across the integration junction (Fig. S2). Clones positives for single crossover mutants were streaked onto *C. difficile* minimal medium (CDMM) supplemented with 5 µg/mL uracil and 2 mg/mL 5-fluoroorotic acid (FOA) to select for plasmid excision. The isolated FOA-resistant colonies were screened by PCR using primers P3 and P4. Double-crossover mutants, in which the mutant allele was successfully integrated yielded products smaller than those seen in WT revertants (1630 bp instead of 2472 bp). In order to restore the *pyrE*^+^ phenotype, plasmid pMTL-YN1 carrying the WT *pyrE* allele was conjugated into the isolated double-crossover mutants. The resulting colonies were restreaked onto non-supplemented CDMM agar to select for uracil prototrophy indicating successful allele exchange.

Successful restoration of the WT *pyrE* allele was confirmed by colony PCR using primers flanking the *pyrE* locus (P5 and P6, Fig. S2).

### *In trans* complementation of the Δ *csiA* mutation

To complement the *csiA* mutation the coding sequence and its expected promoter region were amplified by PCR using primers *csiA* _Fwd_BamHI and *csiA* _Rev_XhoI yielding a 1515 bp fragment. The fragment was then digested with BamHI and XhoI and inserted between the same sites of pMTL-YN1 (Ng *et al*., 2013), yielding pAM38. This plasmid was introduced into *E. coli* HB101 (RP4) and then transferred to strain 630 *erm* Δ*pyrE* Δ *csiA* by conjugation (Heap *et al*., 2007). Following two passages on BHI agar supplemented with 25 µg/mL cefoxitin and 15 µg/mL thiamphenicol, colonies that were noticeably larger (indicative of plasmid integration) were streaked onto onto non-supplemented CDMM to select for uracil prototrophy indicating successful allele exchange. Successful restoration of the WT *pyrE* allele was confirmed by colony PCR using primers flanking the *pyrE* locus (P5 and P6, Fig. S2).

### *SNAP^Cd^* transcriptional fusions

To construct transcriptional *SNAP^Cd^* fusions to the *spo0A* promoter, a 568 bp DNA fragment containing the Spo0A promoter region was PCR-amplified using genomic DNA from strain 630 Δ*erm* and primer pairs CDspo0A_598_Fw/CDspo0A_SNAP_Rev. These fragments were cloned into pFT47 (Pereira *et al*., 2013) to create pMS463 (Table S3). Plasmid pMS463 was transferred to 630Δ*erm*, and congenic Δ*csiA* mutants by conjugation from derivatives of *E. coli* HB101 (RP4) (Table S1).

### SNAP^Cd^ labelling, fluorescence microscopy and image analysis

Samples of 1 ml were withdrawn from SM cultures at the desired times following inoculation, and the cells collected by centrifugation (10 min, 4000xg, at 4°C). The cells were washed with 1ml of phosphate-buffered saline (PBS; 137 mM NaCl, 10 mM Phosphate, 2.7 mM KCl, pH 7.4), and resuspended in 0.1 ml of PBS supplemented with the lipophilic styryl membrane dye *N*-(3-triethylammoniumprpyl)-4-(*p*-diethylaminophenyl-hexatrienyl) pyridinium dibromide (FM4-64, Molecular Probes, Invitrogen; 10 μg.ml^-1^) (Pogliano *et al*., 1999). For SNAP labelling, TMR-Star was added to cells in culture samples inside an anaerobic chamber to a final concentration of 250 nM (New England Biolabs) and the mixture incubated for 30 minutes in the dark. Following labelling, the cells were collected by centrifugation (4000xg for 5 min), washed four times with 1 ml of PBS, and finally suspended in 10-20 μl of PBS. For phase contrast and fluorescence microscopy, cells were mounted on 1.7% agarose coated glass slides and observed on a Leica DM6000B microscope equipped with a phase contrast Uplan F1 100x objective and captured with a CCD Andor Ixon camera (Andor Technologies). Images were acquired and analysed using the Metamorph software suite (version 5.8; Universal Imaging) and adjusted and cropped using Photoshop S4. Statistical analysis was carried out using GraphPad Prism (Version 7.0; GraphPad Software Inc.). The non-parametric Kolmogorov-Smirnov test (KS-test) was applied to compare distributions obtained from quantifications of the SNAP-TMR signal. The P-value is indicated for all comparisons whose differences were found to be statistically significant. Although the results presented are from a single experiment, all experiments involving quantification of a fluorescence signal were performed independently three times and only results that were considered statistically significant by a KS-test in all three experiments were considered to be statistically relevant.

### Preparation of *C. difficile* whole cell extracts and immunoblotting

Whole cell extracts were obtained by withdrawing 20 ml samples from SM or TY cultures of *C. difficile* 6, 8, 10 and 12 hours of after inoculation. The cells were collected by centrifugation (4000xg, for 5 min at 4°C), the cell sediment was washed with phosphate-buffered saline (PBS) and suspended in 1 ml French press buffer (10 mM Tris pH 8.0, 10 mM MgCl2, 0.5 mM EDTA, 0.2 mM NaCl, 10% Glycerol, 1 mM PMSF). The cells were lysed using a French pressure cell (18000 lb/in2). Proteins in the extracts were resolved on 12% SDS-PAGE gels. The anti-σ^F^, anti-σ^E^ antibodies (Saujet *et al*., 2013), anti-Spo0A (Serra *et al*., 2014) and anti-CsiA were used at a 1:1000 dilution, and an anti-rabbit secondary antibody conjugated to horseradish peroxidase (Sigma) was used at a 1:10000 dilution. The monoclonal anti-TcdA primary antibody (Santa Cruz Biotechnology) were used at a 1:1000 dilution, and an anti-mouse secondary antibody conjugated to horseradish peroxidase (Sigma) was used at a 1:2000 dilution. The immunoblots were developed with enhanced chemiluminescence reagents (Amersham Pharmacia Biotech). Images were adjusted and cropped and quantified using ImageJ (http://rsbweb.nih.gov/ij/).

### Detection of Spo0A phosphorylation using Phos-tag acrylamide gels

Phos-tag acrylamide gels were prepared according to the instructions provided (Wako); 10% acrylamide gels were copolymerized with 25 nM Phos-tag acrylamide and 10 nM MnCl2. 20-ml of bacterial cultures were centrifuged at 5,000 *g* at 4°C for 10 min, and the pellets suspended in 1 ml 10 mM Tris pH 8.0. The cells were lysed using a French pressure cell (18000 lb/in2). Samples were stored on ice prior to loading onto Phos-tag acrylamide gels and run at 4°C. Gels were fixed for 10 min in transfer buffer with 10 mM EDTA and then washed for 10 min in transfer buffer without EDTA twice. After transfer to a nitrocellulose membrane, the samples were probed with rabbit polyclonal anti-Spo0A or anti-FliC antibodies at a 1:1000 dilution and an anti-rabbit secondary antibody conjugated to horseradish peroxidase (Sigma) was used at dilution 1:10000. Images were adjusted and cropped and quantified using ImageJ (http://rsbweb.nih.gov/ij/). In order to dephosphorylate Spo0A∼P, samples were incubated at 100°C for 5 min.

### RNA extraction and RNA-sequencing

RNA for RNA-Seq was extracted from two independent biological replicates of WT and *csiA* mutant *C. difficile* strains, after 10 hours of growth in SM, using a RNeasy Kit (Qiagen). Contaminating genomic DNA was depleted by two DNase treatments (Promega), according to manufacturer’s recommendations. DNAse-treated RNA (5 µg) was mRNA enriched using a Ribo-Zero Magnetic Kit (Epicentre). After Illumina sequencing the reads were mapped to *C. difficile 630* genome using Hisat. Statistical analyses were performed with DESeq2. A gene was considered differentially expressed when the fold change was > 2 and the adjusted p value was < 0.01.

### His_6_-CsiA overproduction purification and polyclonal antibody production

A DNA fragment encoding the *csiA* gene was generated by PCR from *C. difficile* 630 Δ*erm* genomic DNA using primers csiA_BamHI_Fw/ csiA_NotI_Rev. The resulting DNA fragment was cut with BamHI and NotI and cloned between the same sites of pETDuet-1 (Novagen) to produce pDM35. Plasmid pDM35 was introduced into BL21 (DE3) cells and the *E. coli* strain was grown in autoinduction medium. The cells were then harvested by centrifugation (4000 x *g*, for 10 min, at 4°C) and the sediment resuspended in lysis buffer (20mM phosphate pH7.4, 1 mM PMSF, 10 mM Imidazole). The suspension was lysed using a French pressure cell (at 18000 lb/in^2^) and the lysate cleared by centrifugation (15000 x *g*, 30 min at at 4°C), and the supernantant was loaded onto a 1 ml Histrap column (Amersham Phamarcia Biotech). The bound protein was eluted with a discontinuous imidazole gradient and the fractions containing His_6_-CsiA were identified by SDS-PAGE. The antibody was produced by Eurogentec (Seraing, Belgium).

## Acknowledgments

This work was supported by the FCT (“Fundação para a Ciência e a Tecnologia”) through program IF (IF/00268/2013/CP1173/CT0006) and award PTDC/BIA-MIC/ to MS, by the European Union Marie Sklodowska Curie 29293/2017 Innovative Training Networks (contract number 642068) to AOH and by awards 5R01AI116895 and 1U01AI124290 from the National Institute of Allergy and Infectious Diseases to SMB. This work was also financially supported by Project LISBOA-01-0145-FEDER-007660 (“Microbiologia Molecular, Estrutural e Celular”) funded by FEDER funds through COMPETE2020 – “Programa Operacional Competitividade e Internacionalização” (POCI) and partially supported by project ONEIDA (LISBOA-01-0145-FEDER-016417) co-funded by FEEI - “Fundos Europeus Estruturais e de Investimento” from “Programa Operacional Regional Lisboa 2020”. ALM was the recipient of a PhD fellowship (PD/BD/105738/2014) within the scope of the PhD program Molecular Biosciences funded by FCT. DM is the recipient of a PhD fellowship (PD/BD/143148/2019) within the scope of the PhD program INTERFACE funded by FCT.

## References

Abecasis, A.B., M. Serrano, R. Alves, L. Quintais, J.B. Pereira-Leal & A.O. Henriques, (2013) A genomic signature and the identification of new sporulation genes. J Bacteriol 195: 2101–2115.

Almeida, A., A.L. Mitchell, M. Boland, S.C. Forster, G.B. Gloor, A. Tarkowska, T.D. Lawley & R.D. Finn, (2019) A new genomic blueprint of the human gut microbiota. Nature 568: 499–504.

Alsaker, K.V., T.R. Spitzer & E.T. Papoutsakis, (2004) Transcriptional analysis of spo0A overexpression in Clostridium acetobutylicum and its effect on the cell’s response to butanol stress. J Bacteriol 186: 1959–1971.

Antunes, A., E. Camiade, M. Monot, E. Courtois, F. Barbut, N.V. Sernova, D.A. Rodionov, Martin-Verstraete & B. Dupuy, (2012) Global transcriptional control by glucose and carbon regulator CcpA in Clostridium difficile. Nucleic Acids Res 40: 10701–10718.

Atarashi, K., T. Tanoue, T. Shima, A. Imaoka, T. Kuwahara, Y. Momose, G. Cheng, S. Yamasaki, T. Saito, Y. Ohba, T. Taniguchi, K. Takeda, S. Hori, Ivanov, II, Y. Umesaki, K. Itoh & K. Honda, (2011) Induction of colonic regulatory T cells by indigenous Clostridium species. Science 331: 337–341.

Ben-Yehuda, S. & R. Losick, (2002) Asymmetric cell division in B. subtilis involves a spiral-like intermediate of the cytokinetic protein FtsZ. Cell 109: 257–266.

Browne, H.P., S.C. Forster, B.O. Anonye, N. Kumar, B.A. Neville, M.D. Stares, D. Goulding & T.D. Lawley, (2016) Culturing of ’unculturable’ human microbiota reveals novel taxa and extensive sporulation. Nature 533: 543–546.

Carroll, K.C. & J.G. Bartlett, (2011) Biology of Clostridium difficile: implications for epidemiology and diagnosis. Annu Rev Microbiol 65: 501–521.

Childress, K.O., A.N. Edwards, K.L. Nawrocki, S.E. Anderson, E.C. Woods & S.M. McBride, (2016) The Phosphotransfer Protein CD1492 Represses Sporulation Initiation in Clostridium difficile. Infect Immun 84: 3434–3444.

Dawson, L.F., E. Valiente, A. Faulds-Pain, E.H. Donahue & B.W. Wren, (2012) Characterisation of Clostridium difficile biofilm formation, a role for Spo0A. PLoS One 7: e50527.

Deakin, L.J., S. Clare, R.P. Fagan, L.F. Dawson, D.J. Pickard, M.R. West, B.W. Wren, N.F. Fairweather, G. Dougan & T.D. Lawley, (2012) The Clostridium difficile spo0A gene is a persistence and transmission factor. Infect Immun 80: 2704–2711.

Dembek, M., S.E. Willing, H.A. Hong, S. Hosseini, P.S. Salgado & S.M. Cutting, (2017) Inducible Expression of spo0A as a Universal Tool for Studying Sporulation in Clostridium difficile. Front Microbiol 8: 1793.

Dineen, S.S., A.C. Villapakkam, J.T. Nordman & A.L. Sonenshein, (2007) Repression of Clostridium difficile toxin gene expression by CodY. Mol Microbiol 66: 206–219.

Driks, A. & P. Eichenberger, (2016) The Spore Coat. Microbiol Spectr 4.

Dworkin, J. & R. Losick, (2005) Developmental commitment in a bacterium. Cell 121: 401–409.

Edwards, A.N., B.R. Anjuwon-Foster & S.M. McBride, (2019) RstA Is a Major Regulator of Clostridioides difficile Toxin Production and Motility. mBio 10.

Edwards, A.N., K.L. Nawrocki & S.M. McBride, (2014) Conserved oligopeptide permeases modulate sporulation initiation in Clostridium difficile. Infect Immun 82: 4276–4291.

Edwards, A.N., R. Tamayo & S.M. McBride, (2016) A novel regulator controls Clostridium difficile sporulation, motility and toxin production. Mol Microbiol 100: 954–971.

Errington, J., (2003) Regulation of endospore formation in Bacillus subtilis. Nat Rev Microbiol 1: 117–126.

Fimlaid, K.A., J.P. Bond, K.C. Schutz, E.E. Putnam, J.M. Leung, T.D. Lawley & A. Shen, (2013) Global analysis of the sporulation pathway of Clostridium difficile. PLoS Genet 9: e1003660.

Fujita, M., J.E. Gonzalez-Pastor & R. Losick, (2005) High- and low-threshold genes in the Spo0A regulon of Bacillus subtilis. J Bacteriol 187: 1357–1368.

Fujita, M. & R. Losick, (2005) Evidence that entry into sporulation in Bacillus subtilis is governed by a gradual increase in the level and activity of the master regulator Spo0A. Genes Dev 19: 2236–2244.

Fujita, M. & Y. Sadaie, (1998) Feedback loops involving Spo0A and AbrB in in vitro transcription of the genes involved in the initiation of sporulation in Bacillus subtilis. J Biochem 124: 98–104.

Galperin, M.Y., S.L. Mekhedov, P. Puigbo, S. Smirnov, Y.I. Wolf & D.J. Rigden, (2012) Genomic determinants of sporulation in Bacilli and Clostridia: towards the minimal set of sporulation-specific genes. Environ Microbiol 14: 2870–2890.

Girinathan, B.P., J. Ou, B. Dupuy & R. Govind, (2018) Pleiotropic roles of Clostridium difficile sin locus. PLoS Pathog 14: e1006940.

Heap, J.T., O.J. Pennington, S.T. Cartman, G.P. Carter & N.P. Minton, (2007) The ClosTron: a universal gene knock-out system for the genus Clostridium. J Microbiol Methods 70: 452–464.

Henriques, A.O. & C.P. Moran, Jr., (2007) Structure, assembly, and function of the spore surface layers. Annu Rev Microbiol 61: 555–588.

Higgins, D. & J. Dworkin, (2012) Recent progress in Bacillus subtilis sporulation. FEMS Microbiol Rev 36: 131–148.

Hussain, H.A., A.P. Roberts & P. Mullany, (2005) Generation of an erythromycin-sensitive derivative of Clostridium difficile strain 630 (630Deltaerm) and demonstration that the conjugative transposon Tn916DeltaE enters the genome of this strain at multiple sites. J Med Microbiol 54: 137–141.

Jiang, M., W. Shao, M. Perego & J.A. Hoch, (2000) Multiple histidine kinases regulate entry into stationary phase and sporulation in Bacillus subtilis. Mol Microbiol 38: 535–542.

Karasawa, T., S. Ikoma, K. Yamakawa & S. Nakamura, (1995) A defined growth medium for Clostridium difficile. Microbiology 141 ( **Pt 2**): 371–375.

Khvorova, A., L. Zhang, M.L. Higgins & P.J. Piggot, (1998) The spoIIE locus is involved in the Spo0A-dependent switch in the location of FtsZ rings in Bacillus subtilis. J Bacteriol 180: 1256–1260.

Koide, A. & J.A. Hoch, (1994) Identification of a second oligopeptide transport system in Bacillus subtilis and determination of its role in sporulation. Mol Microbiol 13: 417–426.

Levin, P.A. & R. Losick, (1996) Transcription factor Spo0A switches the localization of the cell division protein FtsZ from a medial to a bipolar pattern in Bacillus subtilis. Genes Dev 10: 478–488.

Lewis, R.J., J.A. Brannigan, K. Muchova, I. Barak & A.J. Wilkinson, (1999) Phosphorylated aspartate in the structure of a response regulator protein. J Mol Biol 294: 9–15.

McKenney, P.T., A. Driks & P. Eichenberger, (2013) The Bacillus subtilis endospore: assembly and functions of the multilayered coat. Nat Rev Microbiol 11: 33–44.

Molle, V., M. Fujita, S.T. Jensen, P. Eichenberger, J.E. Gonzalez-Pastor, J.S. Liu & R. Losick, (2003) The Spo0A regulon of Bacillus subtilis. Mol Microbiol 50: 1683–1701.

Narula, J., S.N. Devi, M. Fujita & O.A. Igoshin, (2012) Ultrasensitivity of the Bacillus subtilis sporulation decision. Proc Natl Acad Sci U S A 109: E3513–3522.

Nawrocki, K.L., A.N. Edwards, N. Daou, L. Bouillaut & S.M. McBride, (2016) CodY-Dependent Regulation of Sporulation in Clostridium difficile. J Bacteriol 198: 2113–2130.

Ng, Y.K., M. Ehsaan, S. Philip, M.M. Collery, C. Janoir, A. Collignon, S.T. Cartman & N.P. Minton, (2013) Expanding the repertoire of gene tools for precise manipulation of the Clostridium difficile genome: allelic exchange using pyrE alleles. PLoS One 8: e56051.

Paredes-Sabja, D., A. Shen & J.A. Sorg, (2014) Clostridium difficile spore biology: sporulation, germination, and spore structural proteins. Trends Microbiol.

Perego, M., C.F. Higgins, S.R. Pearce, M.P. Gallagher & J.A. Hoch, (1991) The oligopeptide transport system of Bacillus subtilis plays a role in the initiation of sporulation. Mol Microbiol 5: 173–185.

Pereira, F.C., L. Saujet, A.R. Tomé, M. Serrano, M. Monot, E. Couture-Tosi, I. Martin-Verstraete, B. Dupuy & A.O. Henriques, (2013) The spore differentiation pathway in the enteric pathogen Clostridium difficile. PLoS Genet.

Pettit, L.J., H.P. Browne, L. Yu, W.K. Smits, R.P. Fagan, L. Barquist, M.J. Martin, D. Goulding, S.H. Duncan, H.J. Flint, G. Dougan, J.S. Choudhary & T.D. Lawley, (2014) Functional genomics reveals that Clostridium difficile Spo0A coordinates sporulation, virulence and metabolism. BMC Genomics 15: 160.

Piggot, P.J. & D.W. Hilbert, (2004) Sporulation of Bacillus subtilis. Curr Opin Microbiol 7: 579–586.

Pogliano, J., N. Osborne, M.D. Sharp, A. Abanes-De Mello, A. Perez, Y.L. Sun & K. Pogliano, (1999) A vital stain for studying membrane dynamics in bacteria: a novel mechanism controlling septation during Bacillus subtilis sporulation. Mol Microbiol 31: 1149–1159.

Putnam, E.E., A.M. Nock, T.D. Lawley & A. Shen, (2013) SpoIVA and SipL are Clostridium difficile spore morphogenetic proteins. J Bacteriol 195: 1214–1225.

Rosenbusch, K.E., D. Bakker, E.J. Kuijper & W.K. Smits, (2012) C. difficile 630Deltaerm Spo0A regulates sporulation, but does not contribute to toxin production, by direct high-affinity binding to target DNA. PLoS One 7: e48608.

Saujet, L., M. Monot, B. Dupuy, O. Soutourina & I. Martin-Verstraete, (2011) The key sigma factor of transition phase, SigH, controls sporulation, metabolism, and virulence factor expression in Clostridium difficile. J Bacteriol 193: 3186–3196.

Saujet, L., F.C. Pereira, M. Serrano, O. Soutourina, M. Monot, P.V. Shelyakin, M.S. Gelfand, B. Dupuy, A.O. Henriques & I. Martin-Verstraete, (2013) Genome-wide analysis of cell type-specific gene expression during spore formation in Clostridium difficile. PLoS Genet.

Serra, C.R., A.M. Earl, T.M. Barbosa, R. Kolter & A.O. Henriques, (2014) Sporulation during growth in a gut isolate of Bacillus subtilis. J Bacteriol 196: 4184–4196.

Sonenshein, A.L., (2000) Control of sporulation initiation in Bacillus subtilis. Curr Opin Microbiol 3: 561–566.

Steiner, E., A.E. Dago, D.I. Young, J.T. Heap, N.P. Minton, J.A. Hoch & M. Young, (2011) Multiple orphan histidine kinases interact directly with Spo0A to control the initiation of endospore formation in Clostridium acetobutylicum. Mol Microbiol 80: 641–654.

Stewart, G.C., (2015) The Exosporium Layer of Bacterial Spores: a Connection to the Environment and the Infected Host. Microbiol Mol Biol Rev 79: 437–457.

Traag, B.A., A. Pugliese, J.A. Eisen & R. Losick, (2013) Gene conservation among endospore-forming bacteria reveals additional sporulation genes in Bacillus subtilis. J Bacteriol 195: 253–260.

Underwood, S., S. Guan, V. Vijayasubhash, S.D. Baines, L. Graham, R.J. Lewis, M.H. Wilcox & K. Stephenson, (2009) Characterization of the sporulation initiation pathway of Clostridium difficile and its role in toxin production. J Bacteriol 191: 7296–7305.

Vishnoi, M., J. Narula, S.N. Devi, H.A. Dao, O.A. Igoshin & M. Fujita, (2013) Triggering sporulation in Bacillus subtilis with artificial two-component systems reveals the importance of proper Spo0A activation dynamics. Mol Microbiol 90: 181–194.

Wilson, K.H., M.J. Kennedy & F.R. Fekety, (1982) Use of sodium taurocholate to enhance spore recovery on a medium selective for Clostridium difficile. J Clin Microbiol 15: 443–446.

Winkelman, J.T., K.M. Blair & D.B. Kearns, (2009) RemA (YlzA) and RemB (YaaB) regulate extracellular matrix operon expression and biofilm formation in Bacillus subtilis. J Bacteriol 191: 3981–3991.

Winkelman, J.T., A.C. Bree, A.R. Bate, P. Eichenberger, R.L. Gourse & D.B. Kearns, (2013) RemA is a DNA-binding protein that activates biofilm matrix gene expression in Bacillus subtilis. Mol Microbiol 88: 984–997.

Worner, K., H. Szurmant, C. Chiang & J.A. Hoch, (2006) Phosphorylation and functional analysis of the sporulation initiation factor Spo0A from Clostridium botulinum. Mol Microbiol 59: 1000–1012.

Yasugi, M., D. Okuzaki, R. Kuwana, H. Takamatsu, M. Fujita, M.R. Sarker & M. Miyake, (2016) Transcriptional Profile during Deoxycholate-Induced Sporulation in a Clostridium perfringens Isolate Causing Foodborne Illness. Appl Environ Microbiol 82: 2929–2942.

Zhu, D., J.A. Sorg & X. Sun, (2018) Clostridioides difficile Biology: Sporulation, Germination, and Corresponding Therapies for C. difficile Infection. Front Cell Infect Microbiol 8: 29.

