## Supplemental Text for "A regulatory protein that represses sporulation in *Clostridioides difficile*"

**Supplemental Information**

**Supplemental Material and Methods**

**Motility and biofilm formation assays.** Strains were grown in BHI medium to an OD_600_ of 0.5 and 5 µl of culture was spotted into the centre of one-half-concentration BHI plates containing 0.3 % agar, as described (Childress *et al.*, 2016). Swimming diameters were measured every 24 h for a total of 96 h. For the biofilm assay, 1 ml of BHI-S medium containing 0.1 M glucose, 0.1% cysteine and DOC (240μM) was inoculated in a well of a 24-well microplate. Microplates were incubated at 37°C for 24 hours. The biofilm was washed with PBS (Phosphate buffered saline), stained with 1 ml of crystal violet (0.2%) followed by two washes with PBS. The OD600nm was measured after resuspension of the cells in methanol/acetone using non-inoculated medium as a negative control.

**Supplemental Results**

**Impact of *csiA* deletion on motility, biofilm formation and toxin production.** To determine whether the *csiA* mutation had an impact on toxin production, biofilm formation or motility we test the different phenotypes. We first tested motility on semi-solid BHI plates. The swimming motility of the wild-type and the *csiA* mutant on soft agar medium did not differ significantly (Fig. S5A). Since σ^D^ is required for the expression of *fliC* (coding for flagellin) and other motility genes (Dingle *et al.*, 2011), a *sigD* mutant was used as negative control for the experiment (the cells of a *sigD* mutant are not motile). We also tested the ability of these strains to form biofilm in the presence of glucose and DOC, using the cristal violet assay (Poquet *et al.*, 2018). The *csiA* mutant has no effect in the ability of inducing biofilm formation in the presence of glucose and DOC after 24 hours of growth (Fig. S5B). Finally, we used western blot to investigate the levels of toxin (TcdA) inside the cells. The wild-type and the mutant strain were grown in TY, samples were collected 14 hours after inoculation, and the presence of TcdA was monitored in the cell fractions (Fig. S5C). We found similar TcdA levels in the wild type and *csiA* mutant strains.

**Supplemental figures legends**

**Figure S1.** ***csiA* is highly conserved.** Neighbourhood analysis of genes surrounding *csiA* in the genomes of the indicated sporulating bacteria. Named genes are colour coded to indicate conservation in the various genomes. Grey genes are not conserved with other genomes in the *csiA* neighbourhood.

**Figure S2.** **Construction of the** Δ***csiA* in-frame deletion mutant using allele-coupled exchange (ACE).** **A:** PCR fragments A, B and C were obtained with primers pairs P2/P3, P1/P4 and P3/P4, respectively. PCR fragments D and E were obtained with primers P5/P6. Note that following the initial single reciprocal cross-over, only the integration represented first (of the two possible orientations represented) was obtained. The shorter *csiA* gene (in yellow) represents the allele with an in-frame deletion of codons 5 to 291 of the 293-codons-long open reading frame. **B:** PCR analysis of the parental 630Δ*erm* strain in comparison with the *csiA* deletion mutant (Δ*csiA*) and the *csiA* in trans complementation strain (*csiA*^C^, with a copy of the *csiA* gene at the *pyrE* locus). PCR analysis was performed using the primer pair P3/P4 (for *csiA* locus verification) and P5/P6 (for *pyrE* locus verification). The position and sizes of the expected products is shown on the right side of the panel (red arrows). The position on molecular size markers (M, in bp) is shown on the left side of the panel.

**Figure S3. The** Δ***csiA* mutant shows increased sporulation in SM. A:** Samples of an SM liquid culture of the wild type strain (WT, 630 Δ*erm*), the Δ*csiA* mutant and the complementation strain (*csiA*^C^) were collected at 14 hours after inoculation, stained with the membrane dye FM4-64 and examined by phase contrast and fluorescent microscopy. The numbers in the panels are the percentage of sporulating cells. Data shown are from one representative experiment in which at least 100 cells were analysed for each strain. Scale bar, 1 µm. **B:** The optical density of cultures of the WT and the Δ*csiA* strains was measured at 600nm in intervals of 2 hours until 20 hours hour after inoculation. The data represent the mean ± standard deviation (SD) of three independent experiments.

**Figure S4. The** Δ***csiA* phenotype in 70:30 sporulation medium.	A:** Cells were grown in 70:30 sporulation medium plates and the titer of heat resistant spores and total viable cells measured 12 hours following inoculation. The data in the graph represent the mean of two independent experiments. **B:** Quantitative analysis of the fluorescence intensity (Fl.) of *C. difficile* cells carrying a P*_spoOA_*-*SNAP^Cd^* transcriptional fusion in the WT and congenic Δ*csiA* mutant. Only cells with no signs of sporulation were quantified. Cells that The cells were collected after 10 h of growth in liquid SM (left) or in plates of 70:30 sporulation medium (right) , stained with TMR-Star, and examined by fluorescence microscopy to monitor SNAP production. Data shown are from one experiment, representative of at least three independent experiments. The numbers in the legend represent the mean and the SD of fluorescence intensity.

**Figure S5.** **Disruption of the** Δ***csiA* gene has no effect on motility, biofilm formation or toxin production.** **A:** Motility assays of WT, Δ*csiA* and Δ*sigD* (negative control) strains in one-half-concentration BHI medium fortified with 0.3 % agar. Swimming diameters were measured every 24h for a total of 96h, as indicated. A two-tailed Student *t* test was performed to determine statistically significant differences. **B:** Mean values of the OD600nm measured after crystal violet staining of the mass of biofilm obtained after 24 h in presence of DOC 240 µm. One-way ANOVA was performed to determine statistically significant differences. The data in A and B represent the mean ± SD of three independent experiments performed in triplicate. **C:** Samples were collected from the WT, Δ*csiA* and Δ*tcdAB* mutant grown in TY for 14 hours. Extracts were prepared and proteins (15 µg) resolved by SDS-PAGE and subjected to immunobloting using an anti-TcdA antibody**.** In the bottom panel the same extracts were loaded in a SDS-PAGE and subjected Comassie blue staining, as a loading control. The position of molecular weight markers (in kDa) is indicated on the left side of the panels.

**Supplemental Tables**

**Table S1. Bacterial strains used in this study.**

| Strain | Genotype and phenotype | Origin/Construction |
| --- | --- | --- |
| *E. coli* |  |  |
| DH5α | *fhuA2 lac(del)U169 phoA glnV44 Φ80' lacZ(del)M15 gyrA96 recA1 relA1 endA1 thi-1 hsdR17* | Laboratory Stock |
| HB101 (RP4) | *supE*44 *aa*14 *galK*2 *lacY*1 ∆ (*gpt-proA*) 62 *rpsL*20 (Str^R^)*xyl-5 mtl-1 recA*13 ∆ (*mcrC-mrr*) *hsdS*_B_(r_B_^-^m_B_) RP4 (Tra^+^ IncP Ap^R^ Km^R^ Tc^R^) | Laboratory Stock |
| *C. difficile* |  |  |
| AHCD591 | 630∆*erm tcdAB*::*ermB* | (Kuehne *et al.*, 2010) |
| AHCD597 | 630Δ*erm*P*_sigE_*-*SNAP* | (Pereira *et al.*, 2013) |
| AHCD598 | 630Δ*erm* P*_sigF_*-*SNAP* | “ |
| AHCD602 | 630Δ*erm* P*_gpr_*-*SNAP* | “ |
| AHCD603 | 630Δ*erm* P*_spoIIIA_*-*SNAP* | “ |
| AHCD607 | 630Δ*erm* P*_spo0A_*-*SNAP* | This work |
| AHCD683 | 630∆*erm sigD*::*ermB* | (Aubry *et al.*, 2012) |
| AHCD772 | *630ΔermΔpyrE* | (Ng *et al.*, 2013) |
| AHCD1014 | *630ΔermΔpyrEΔcsiA* | This work |
| AHCD1034 | 630Δ*erm* P*_csiA_*-*SNAP* | “ |
| AHCD1055 | *630Δerm ΔcsiA pyrE:: csiA* | “ |
| AHCD1077 | 630Δ*erm* Δ*csiA* | “ |
| AHCD1131 | 630Δ*erm* Δ*csiA* P*_gpr_*-*SNAP* | “ |
| AHCD1132 | 630Δ*erm* Δ*csiA* P*_spoIIIA_*-*SNAP* | “ |
| AHCD1142 | 630Δ*erm* Δ*csiA* P*_spo0A_*-*SNAP* | “ |
| AHCD1143 | 630Δ*erm* Δ*csiA* P*_sigF_*-*SNAP* | “ |
| AHCD1146 | 630Δ*erm* Δ*csiA* P*_sigE_*-*SNAP* | “ |
| AHCD1190 | *630Δerm* | “ |
| AHCD1248 | 630Δ*erm* Δ*spo0A* P*_spo0A_*-*SNAP* | “ |
| AHCD1263 | 630Δ*erm* Δ*csiA* P*_tet_*-*gusA* | “ |
| AHCD1272 | 630Δ*erm* P*_tet_*-*gusA* | “ |
| AHCD1274 | 630Δ*erm* Δ*csiA* P*_tet_*-*spo0A* | “ |
| AHCD1279 | 630Δ*erm* P*_tet_*-*spo0A* | “ |
| AHCD1450 | 630Δ*erm* P*_tet_*-*spo0A* P*_sigE_*-*SNAP* | “ |
| AHCD1455 | 630Δ*erm* P*_tet_*-*spo0A* P*_sigF_*-*SNAP* | “ |

**Table S2. Oligonucleotide used in this study.**

| Name | Sequence (5’ to 3’) |
| --- | --- |
| csiA_AscI_Fwd | ACAGGCGCGCCATAGAAGATTTTCTGTAGGAG |
| csiA_LHA_Rev | ATTCTTACTCTATACTTATAGCCATTAATTAAACCTCC |
| csiA_RHA_Fwd | ATAGAGTAAGAATACAATATTAGAAGGGG |
| csiA_SbfI_Rev | CTTCCTGCAGGCTCTTTGAAACAGTCCCTAC |
| csiA_Fwd_BamHI | GAAGGATCCGTAGATACATTAGATGG |
| csiA_Rev_XhoI | CAACTCGAGCTTTTCCTGCACCTGATGG |
| P1 | TTCTTTCTATTCAGCACTGTTATGC |
| P2 | CATCAAGAAGAGCGACTTCG |
| P3 | GCAATAGATGGAAGATTCGACC |
| P4 | GCTACACATACAGGCTTTTCC |
| P5 | CAATAATTTTATAACATTAACATGG |
| P6 | ATTTACATTTTTTAAGTAACAC |
| csiA_Fw | CGGAATTCGATATAAAAATAAATCTTACC |
| csiA_Rev | CACAATCTTTATCCATTAATTAAACCTCCAAAATACC |
| SNAP_SOE_Fw | ATGGATAAAGATTGTGAAATGAAGAGAACC |
| XhoI_SNAP_Rev | CCGCTCGAGTTACCCAAGTCCTGGTTTCCCCAAACG |
| CDspo0A_598D | GCTAGGATCCTTAATGGGTAATTTC |
| CDspo0A_SNAP_Rev | CCCCATTAAAAACTCGAGTCTTATTACAGA |
| *csiA* 970D | GGAATTCATATGTGAAGGTATTTTGGAGG |
| *csiA* 1920R | ATAAGAATGCGGCCGCCACAAACATTTAAGCATTCCCC |
| *csiA*_BamHI_Fw | CGGGATCCGATGGCTATAAGTATGACTGG |
| *csiA*_NotI_Rev | GCGGCCGCTTACTCTATATTTTGTATTTGTTC |

Underlined sequences represent introduced restriction sites.

**Table S3. Plasmids used in this study.**

| Plasmid | Relevant genotype | Origin |
| --- | --- | --- |
| pFT47 | pMTL84121-*SNAP^Cd^* (Cm^R^/Tm^R^) | (Pereira *et al.*, 2013) |
| pFT48 | pFT47- P*_sigF_*-*SNAP* (Cm^R^/Tm^R^) | “ |
| pFT49 | pFT47-P*_sigE_*-*SNAP* (Cm^R^/Tm^R^) | “ |
| pFT53 | pFT47- P*_gpr_*-*SNAP* (Cm^R^/Tm^R^) | “ |
| pFT54 | pFT47- P*_spoIIIA_*-*SNAP* (Cm^R^/Tm^R^) | “ |
| pMTL84121 | *Clostridia* modular plasmid;*catP* (Cm^R^/Tm^R^) | (Heap *et al.*, 2007) |
| pMTL-YN3 | Plasmid for mutant construction through ACE methodology (Cm^R^/Tm^R^) | (Ng *et al.*, 2013) |
| pMTL-YN1 | Plasmid for Δ*pyrE* reversion through ACE methodology (Cm^R^/Tm^R^) | (Ng *et al.*, 2013) |
| pMTL-YN1C | Plasmid for *in trans* complementation of mutants (Cm^R^/Tm^R^) | (Ng *et al.*, 2013) |
| pMLD138 | pMTL-YN3-P*_tet_*-spo0A | (Dembek *et al.*, 2017) |
| pAM25 | pMTL84121 - P*_tet_* | This work |
| pAM37 | pMTL-YN3 containing homology regions for Δ*csiA* construction (Cm^R^/Tm^R^) | “ |
| pAM38 | pMTL-YN1C containing *csiA* (Cm^R^/Tm^R^) | “ |
| pDM35 | pETDuet-1-*csiA* (Amp^R^) | “ |
| pMS463 | pFT47- P*_spo0A_*-*SNAP* (Cm^R^/Tm^R^) | “ |
