## Supplemental Table 3 for "A regulatory protein that represses sporulation in *Clostridioides difficile*"

**T10**
**Adjusted p-value <0.01**

| | | | | | | | | | | | | | |
|---|---|---|---|---|---|---|---|---|---|---|---|---|---|
| **>4 fold** | | | | | | | | | | | | **>4 fold** | |
| GeneID | Base mean | log2 (FC) | StdErr | Wald-Stats | p-value | p-adj | Gene ID | Counts-y1 | Counts-y2 | Counts-wt1 | Counts-wt2 | GeneID | |
| CD630_25890 | 742,2377 | 8,814003 | 0,554696 | 15,88978 | 7,46E-57 | 2,88E-53 | CD630_25 | 0,900608 | 0 | 1451,091 | 1516,959068 | CD630_25890 | csiA |
| CD630_22150 | 393,896 | -5,297177 | 0,437403 | -12,11053 | 9,29E-34 | 1,79E-30 | CD630_22 | 835,7638 | 710,8656 | 16,957658 | 11,99704008 | CD630_22150 | sinR2 |
| CD630_23990 | 1754,292 | -4,359196 | 0,385452 | -11,30932 | 1,18E-29 | 1,52E-26 | CD630_23 | 4395,865 | 2354,944 | 128,39369 | 137,9659609 | CD630_23990 | conserved hypotetical protein K CotJB1? |
| CD630_25990 | 520,8707 | -4,2162 | 0,383798 | -10,98546 | 4,49E-28 | 4,33E-25 | CD630_25 | 1066,319 | 928,0566 | 48,45045 | 40,65663583 | CD630_25990 | transcriptional regulator G-ClnR activates this (Shonna) |
| CD630_22140 | 642,0691 | -4,512416 | 0,419085 | -10,7673 | 4,91E-27 | 3,79E-24 | CD630_22 | 1451,779 | 1037,619 | 24,225225 | 54,6531826 | CD630_22140 | sinR1 |
| CD630_21152 | 77,26572 | -4,714773 | 0,605147 | -7,791119 | 6,64E-15 | 7,76E-13 | CD630_21 | 145,8984 | 159,832 | 0 | 3,332511134 | CD630_21152 | hypothetical protein |
| **>3 fold** | | | | | | | | | | | | **>3fold** | |
| GeneID | Base mean | log2 (FC) | StdErr | Wald-Stats | p-value | p-adj | Gene ID | Counts-y1 | Counts-y2 | Counts-wt1 | Counts-wt2 | **GeneID** | |
| CD630_14860 | 9622,313 | -3,362623 | 0,336005 | -10,00765 | 1,41E-23 | 9,07E-21 | CD630_14 | 15613,83 | 19808,85 | 1717,5685 | 1349,000507 | CD630_14860 | Putative ribosome recycling factor G |
| CD630_02140 | 895,2066 | -3,925211 | 0,395009 | -9,93701 | 2,87E-23 | 1,58E-20 | CD630_02 | 2205,588 | 1192,295 | 89,633333 | 93,31031175 | CD630_02140 | hypothetical protein F/G and E |
| CD630_26871 | 13714,77 | -3,577978 | 0,377022 | -9,490096 | 2,31E-21 | 1,11E-18 | CD630_26 | 25996,04 | 25207,69 | 2444,3252 | 1211,034546 | CD630_26871 | hypothetical protein F/G |
| CD630_35800 | 5113,604 | -3,396744 | 0,362004 | -9,383165 | 6,40E-21 | 2,74E-18 | CD630_35 | 10667,7 | 8221,679 | 973,85405 | 591,1874752 | CD630_35800 | hypothetical protein K and spoIIID-spore coat? |
| CD630_05430 | 1178,604 | -3,22409 | 0,354386 | -9,097682 | 9,23E-20 | 3,56E-17 | CD630_05 | 2298,35 | 2006,278 | 239,82973 | 169,9580678 | CD630_05430 | hypothetical protein G |
| CD630_12900 | 594,8953 | -3,609153 | 0,400431 | -9,013167 | 2,00E-19 | 6,43E-17 | CD630_12 | 1426,562 | 801,0933 | 79,943243 | 71,98224049 | CD630_12900 | SAPS - G |
| CD630_21501 | 1424,791 | -3,872208 | 0,429461 | -9,01644 | 1,94E-19 | 6,43E-17 | CD630_21 | 1843,544 | 3559,484 | 203,49189 | 92,64380952 | CD630_21501 | hypothetical protein |
| CD630_21410 | 2683,488 | -3,025213 | 0,337194 | -8,971742 | 2,92E-19 | 8,66E-17 | CD630_21 | 5490,103 | 4176,899 | 518,41982 | 548,5313326 | CD630_21410 | serine-type D-Ala-D-Ala carboxypeptidase F |
| CD630_23150 | 594,4266 | -3,463466 | 0,391056 | -8,8567 | 8,24E-19 | 2,27E-16 | CD630_23 | 1169,889 | 1034,396 | 111,43604 | 61,98470709 | CD630_23150 | hypothetical protein G |
| CD630_21120 | 11299,03 | -3,029037 | 0,342325 | -8,848421 | 8,88E-19 | 2,28E-16 | CD630_21 | 16350,53 | 24376,31 | 2151,2 | 2318,094745 | CD630_21120 | hypothetical protein G |
| CD630_28090 | 19464,41 | -3,417222 | 0,39121 | -8,735001 | 2,44E-18 | 5,22E-16 | CD630_28 | 45950,8 | 26191,82 | 3662,854 | 2052,160356 | CD630_28090 | hypothetical protein G |
| CD630_24430 | 654,3255 | -3,828389 | 0,443969 | -8,623105 | 6,52E-18 | 1,32E-15 | CD630_24 | 689,8654 | 1791,02 | 75,098198 | 61,31820486 | CD630_24430 | hypothetical protein E |
| CD630_10631 | 3448,476 | -3,02777 | 0,36924 | -8,200015 | 2,40E-16 | 4,03E-14 | CD630_10 | 6224,999 | 6222,49 | 874,53063 | 471,8835766 | CD630_10631 | hypothetical protein K |
| CD630_17260 | 7795,376 | -3,174629 | 0,390852 | -8,122323 | 4,57E-16 | 7,06E-14 | CD630_17 | 15049,15 | 13425,89 | 1865,3423 | 841,1258102 | CD630_17260 | hypothetical protein E |
| CD630_10280 | 993,3143 | -3,430317 | 0,43196 | -7,941276 | 2,00E-15 | 2,66E-13 | CD630_10 | 1849,848 | 1840,001 | 210,75946 | 72,64874272 | CD630_10280 | diguanylate cyclase G GGDEF domain |
| CD630_02130 | 1078,832 | -3,044146 | 0,385342 | -7,899848 | 2,79E-15 | 3,47E-13 | CD630_02 | 2383,908 | 1517,759 | 254,36486 | 159,2940322 | CD630_02130 | cotF E |
| CD630_32510 | 6993,698 | -3,129061 | 0,409523 | -7,640736 | 2,16E-14 | 2,31E-12 | CD630_32 | 12039,32 | 13470,35 | 1773,2865 | 691,8293114 | CD630_32510 | dehydrogenase E |
| CD630_17880 | 688,6987 | -3,078522 | 0,414255 | -7,431474 | 1,07E-13 | 1,04E-11 | CD630_17 | 1331,098 | 1168,449 | 179,26667 | 75,98125385 | CD630_17880 | hypothetical protein G |
| CD630_23950 | 486,486 | -3,155362 | 0,443962 | -7,107285 | 1,18E-12 | 8,61E-11 | CD630_23 | 533,1597 | 1247,72 | 101,74595 | 63,31771154 | CD630_23950 | hypothetical protein E |
| CD630_31510 | 166,071 | -3,304611 | 0,470207 | -7,027989 | 2,10E-12 | 1,39E-10 | CD630_31 | 223,3507 | 391,2016 | 29,07027 | 20,66156903 | CD630_31510 | hypothetical protein E |
| CD630_20550 | 249,3705 | -3,423286 | 0,492975 | -6,944135 | 3,81E-12 | 2,37E-10 | CD630_20 | 434,0928 | 493,6745 | 55,718018 | 13,99654676 | CD630_20550 | hypothetical protein E/K |
| CD630_11930 | 4175,981 | -3,04417 | 0,444086 | -6,854905 | 7,14E-12 | 4,33E-10 | CD630_11 | 5698,144 | 9493,89 | 1131,318 | 380,5727715 | CD630_11930 | SpoIIIAB E |
| CD630_26860 | 798,0271 | -3,109758 | 0,456333 | -6,814674 | 9,45E-12 | 5,36E-10 | CD630_26 | 1371,625 | 1542,894 | 215,6045 | 61,98470709 | CD630_26860 | membrane protein F |
| CD630_19400 | 9059,59 | -3,077314 | 0,465039 | -6,61733 | 3,66E-11 | 1,88E-09 | CD630_19 | 12253,67 | 20856,78 | 2446,7477 | 681,1652758 | CD630_19400 | membrane protein (DUF3866 superfamily) E |
| CD630_31501 | 103,7792 | -3,194542 | 0,490113 | -6,517975 | 7,13E-11 | 3,51E-09 | CD630_31 | 173,8173 | 210,1017 | 14,535135 | 16,66255567 | CD630_31501 | hypothetical protein E |
| CD630_21430 | 153,2186 | -3,048391 | 0,471204 | -6,469364 | 9,84E-11 | 4,69E-09 | CD630_21 | 246,7665 | 309,3522 | 38,76036 | 17,99556012 | CD630_21430 | HTH-type transcriptional regulator |
| CD630_19060 | 26,87575 | -3,028447 | 0,693738 | -4,365401 | 1,27E-05 | 0,000149 | CD630_19 | 31,52126 | 70,89321 | 2,4225225 | 2,666008907 | CD630_19060 | hypothetical protein |
| **>2 fold** | | | | | | | | | | | | **>2 fold** | |
| GeneID | Base mean | log2 (FC) | StdErr | Wald-Stats | p-value | p-adj | Gene ID | Counts-y1 | Counts-y2 | Counts-wt1 | Counts-wt2 | | |
| CD630_12300 | 20680,33 | -2,869729 | 0,324574 | -8,841513 | 9,44E-19 | 2,28E-16 | CD630_12 | 39712,29 | 33837,97 | 4244,2594 | 4926,78446 | CD630_12300 | sigK E |
| CD630_34640 | 29339,44 | -2,803163 | 0,320076 | -8,757802 | 1,99E-18 | 4,52E-16 | CD630_34 | 50247,6 | 53492,15 | 7267,5675 | 6350,433217 | CD630_34640 | GerS E |
| CD630_29670 | 2620,63 | -2,763155 | 0,334397 | -8,263098 | 1,42E-16 | 2,74E-14 | CD630_29 | 5184,798 | 4060,247 | 605,63063 | 631,844111 | CD630_29670 | spoVFB K |
| CD630_13540 | 791,8847 | -2,976955 | 0,361686 | -8,230778 | 1,86E-16 | 3,42E-14 | CD630_13 | 1598,578 | 1251,587 | 138,08378 | 179,289099 | CD630_13540 | hypothetical protein G |
| CD630_11330 | 9609,206 | -2,787066 | 0,339158 | -8,217596 | 2,08E-16 | 3,64E-14 | CD630_11 | 17669,02 | 16304,79 | 2638,127 | 1824,883097 | CD630_11330 | hypothetical protein K |
| CD630_32490 | 167579,3 | -2,906712 | 0,355665 | -8,172623 | 3,02E-16 | 4,85E-14 | CD630_32 | 343540,3 | 255558,4 | 27725,77 | 43492,60281 | CD630_32490 | sspB G |
| CD630_14300 | 9383,753 | -2,906269 | 0,359867 | -8,075951 | 6,70E-16 | 9,93E-14 | CD630_14 | 21308,37 | 12249,06 | 1930,7504 | 2046,828338 | CD630_14300 | pdaA G |
| CD630_16310 | 2197,314 | -2,620308 | 0,326755 | -8,019174 | 1,06E-15 | 1,52E-13 | CD630_16 | 3932,052 | 3715,449 | 576,56036 | 565,1938883 | CD630_16310 | sodA G |
| CD630_29680 | 2104,228 | -2,630319 | 0,330091 | -7,968475 | 1,61E-15 | 2,21E-13 | CD630_29 | 3916,742 | 3415,119 | 557,18018 | 527,8697636 | CD630_29680 | dpaA K |
| CD630_19270 | 934,7499 | -2,856239 | 0,361344 | -7,904477 | 2,69E-15 | 3,46E-13 | CD630_19 | 1964,225 | 1364,372 | 220,44955 | 189,9531346 | CD630_19270 | ytlC ABC-transporter K |
| CD630_14330 | 17127,92 | -2,745794 | 0,349562 | -7,854963 | 4,00E-15 | 4,82E-13 | CD630_14 | 37205,9 | 23194,32 | 3897,8387 | 4213,627078 | CD630_14330 | cotE K |
| CD630_30240 | 1413,497 | -2,870177 | 0,372563 | -7,703869 | 1,32E-14 | 1,50E-12 | CD630_30 | 3207,063 | 1838,068 | 314,92793 | 293,927482 | CD630_30240 | hypothetical protein |
| CD630_10650 | 18787,27 | -2,451948 | 0,318604 | -7,695898 | 1,41E-14 | 1,55E-12 | CD630_10 | 32804,63 | 31459,18 | 5179,3531 | 5705,925563 | CD630_10650 | hypothetical protein K |
| CD630_21420 | 814,0315 | -2,637333 | 0,349973 | -7,535809 | 4,85E-14 | 5,06E-12 | CD630_21 | 1490,505 | 1350,193 | 230,13964 | 185,287619 | CD630_21420 | transporter |
| CD630_08960 | 1426,276 | -2,643703 | 0,354057 | -7,466886 | 8,21E-14 | 8,33E-12 | CD630_08 | 2952,191 | 2032,057 | 387,6036 | 333,2511134 | CD630_08960 | hypothetical protein K |
| CD630_00160 | 5194,459 | -2,662615 | 0,360283 | -7,390342 | 1,46E-13 | 1,32E-11 | CD630_00 | 6671,701 | 11527,24 | 1264,5568 | 1314,342391 | CD630_00160 | dnaX E |
| CD630_23750 | 42988,53 | -2,98851 | 0,404197 | -7,393702 | 1,43E-13 | 1,32E-11 | CD630_23 | 87879,48 | 67369,17 | 11787,995 | 4917,453429 | CD630_23750 | hypothetical protein F/G |
| CD630_32980 | 10860,1 | -2,53465 | 0,342983 | -7,390024 | 1,47E-13 | 1,32E-11 | CD630_32 | 19111,79 | 18417,41 | 3561,1081 | 2350,086852 | CD630_32980 | ATP/GTP-binding protein E |
| CD630_10660 | 3212,055 | -2,499543 | 0,338588 | -7,382247 | 1,56E-13 | 1,36E-11 | CD630_10 | 4642,632 | 6415,836 | 886,64324 | 903,1105173 | CD630_10660 | hypothetical protein E |
| CD630_14020 | 1267,386 | -2,823889 | 0,382827 | -7,376414 | 1,63E-13 | 1,39E-11 | CD630_14 | 2071,397 | 2434,215 | 370,64594 | 193,2856458 | CD630_14020 | glpQ E |
| CD630_35670 | 86019,72 | -2,371246 | 0,322222 | -7,359041 | 1,85E-13 | 1,55E-11 | CD630_35 | 149099,2 | 142670 | 28660,864 | 23648,83201 | CD630_35670 | SipL E |
| CD630_24351 | 1217,934 | -2,71313 | 0,371159 | -7,309885 | 2,67E-13 | 2,19E-11 | CD630_24 | 2360,492 | 1925,073 | 368,22342 | 217,9462282 | CD630_24351 | hypothetical protein (ykz-like) Spo0A |
| CD630_33500 | 4170,258 | -2,874745 | 0,394439 | -7,288179 | 3,14E-13 | 2,52E-11 | CD630_33 | 10066,99 | 4855,54 | 726,75675 | 1031,745447 | CD630_33500 | family 2 glycosyl transferase E/K |
| CD630_28410 | 3111,832 | -2,368394 | 0,328958 | -7,199682 | 6,04E-13 | 4,75E-11 | CD630_28 | 5798,111 | 4759,512 | 939,93874 | 949,7656732 | CD630_28410 | amidohydrolase G |
| CD630_03110 | 78187,15 | -2,369678 | 0,33128 | -7,153098 | 8,48E-13 | 6,54E-11 | CD630_03 | 146914,3 | 118437,4 | 21015,383 | 26381,49114 | CD630_03110 | putative coat assembly protein E |
| CD630_28160 | 3572,195 | -2,872964 | 0,402637 | -7,135377 | 9,65E-13 | 7,30E-11 | CD630_28 | 6624,869 | 6156,109 | 1065,9099 | 441,8909764 | CD630_28160 | hypothetical protein E |
| CD630_07930 | 334,9444 | -2,938587 | 0,413129 | -7,113 | 1,14E-12 | 8,42E-11 | CD630_07 | 660,1453 | 542,0108 | 89,633333 | 47,98816033 | CD630_07930 | putative membrane protein G |
| CD630_24310 | 4233,107 | -2,592759 | 0,36593 | -7,085399 | 1,39E-12 | 9,72E-11 | CD630_24 | 9465,385 | 5275,744 | 1128,8955 | 1062,404549 | CD630_24310 | nitrite/sulfite reductase G |
| CD630_19040 | 932,2796 | -2,47184 | 0,34922 | -7,078166 | 1,46E-12 | 1,01E-10 | CD630_19 | 1418,457 | 1783,286 | 283,43513 | 243,939815 | CD630_19040 | ABC transporter permease K |
| CD630_24000 | 2836,408 | -2,448441 | 0,347678 | -7,042275 | 1,89E-12 | 1,28E-10 | CD630_24 | 5760,286 | 3959,064 | 881,7982 | 744,4829873 | CD630_24000 | spore coat peptide assembly protein CotJB2 K |
| CD630_12340 | 3118,927 | -2,341405 | 0,333753 | -7,015386 | 2,29E-12 | 1,50E-10 | CD630_12 | 4619,216 | 5935,695 | 915,71351 | 1005,085358 | CD630_12340 | Recombination directionality factor (skin excision) E |
| CD630_25980 | 1416,307 | -2,473318 | 0,353977 | -6,987227 | 2,80E-12 | 1,80E-10 | CD630_25 | 2838,715 | 2028,19 | 445,74414 | 352,579678 | CD630_25980 | oligosaccharide deacetylase G |
| CD630_30320 | 1481,673 | -2,355464 | 0,338696 | -6,954504 | 3,54E-12 | 2,24E-10 | CD630_30 | 2790,983 | 2231,847 | 469,96937 | 433,8929496 | CD630_30320 | pyridoxal phosphate-dependent transferase K |
| CD630_15110 | 43496,67 | -2,287961 | 0,333819 | -6,853903 | 7,19E-12 | 4,33E-10 | CD630_15 | 75482,62 | 70804,92 | 16134 | 11565,14664 | CD630_15110 | cotB E |
| CD630_09020 | 1636,358 | -2,587071 | 0,37786 | -6,846637 | 7,56E-12 | 4,49E-10 | CD630_09 | 3169,238 | 2527,021 | 549,91261 | 299,2594998 | CD630_09020 | cation efflux protein K |
| CD630_18840 | 5428,798 | -2,297262 | 0,335849 | -6,840155 | 7,91E-12 | 4,62E-10 | CD630_18 | 8987,162 | 9297,967 | 1436,5559 | 1993,50816 | CD630_18840 | hypothetical protein E |
| CD630_11680 | 7795,605 | -2,469875 | 0,36129 | -6,836272 | 8,13E-12 | 4,68E-10 | CD630_11 | 13137,16 | 13665,63 | 2798,0135 | 1581,609784 | CD630_11680 | YlbJ-like protein/spore cortex formation E |
| CD630_05710 | 5744,36 | -2,211921 | 0,327411 | -6,755795 | 1,42E-11 | 7,94E-10 | CD630_05 | 9884,168 | 9249,63 | 2129,3973 | 1714,243727 | CD630_05710 | hypothetical protein Spo0A |
| CD630_32480 | 8660,042 | -2,290529 | 0,341755 | -6,702257 | 2,05E-11 | 1,13E-09 | CD630_32 | 14167,46 | 14982,31 | 3284,9405 | 2205,455868 | CD630_32480 | polysaccharide deacetylase E |
| CD630_15430 | 4088,598 | -2,281584 | 0,342183 | -6,667739 | 2,60E-11 | 1,41E-09 | CD630_15 | 8091,058 | 5661,145 | 1218,5288 | 1383,658623 | CD630_15430 | FMN-dependent NADH-azoreductase G |
| CD630_34650 | 7949,406 | -2,129524 | 0,319504 | -6,665099 | 2,64E-11 | 1,42E-09 | CD630_34 | 12594,1 | 13600,54 | 2860,9991 | 2741,990161 | CD630_34650 | hypothetical protein E |
| CD630_34990 | 27666,14 | -2,117466 | 0,317808 | -6,662719 | 2,69E-11 | 1,42E-09 | CD630_34 | 47650,24 | 43363,44 | 9777,3009 | 9873,563987 | CD630_34990 | SpoVT G |
| CD630_26570 | 877,2733 | -2,31436 | 0,348633 | -6,638378 | 3,17E-11 | 1,65E-09 | CD630_26 | 1315,788 | 1648,589 | 256,78739 | 287,928962 | CD630_26570 | hypothetical protein SpoVD operon Spo0A |
| CD630_21840 | 1806,296 | -2,187269 | 0,331922 | -6,589702 | 4,41E-11 | 2,24E-09 | CD630_21 | 2809,895 | 3190,194 | 649,23603 | 575,8579239 | CD630_21840 | Putative N-acetylmuramoyl-L-alanine amidase K |
| CD630_10670 | 52616,34 | -2,094633 | 0,318793 | -6,570514 | 5,01E-11 | 2,51E-09 | CD630_10 | 91483,71 | 81110,86 | 19421,363 | 18449,44814 | CD630_10670 | CdeC K |
| CD630_21590 | 480,8561 | -2,769941 | 0,425058 | -6,516613 | 7,19E-11 | 3,51E-09 | CD630_21 | 734,8957 | 970,5925 | 152,61892 | 65,31721822 | CD630_21590 | putative membrane protein |
| CD630_34570 | 4990,15 | -2,110668 | 0,324168 | -6,511024 | 7,46E-11 | 3,60E-09 | CD630_34 | 8598,1 | 7810,498 | 1894,4126 | 1657,591038 | CD630_34570 | hypothetical protein E |
| CD630_01060 | 6406,289 | -2,124442 | 0,331179 | -6,414779 | 1,41E-10 | 6,63E-09 | CD630_01 | 10860,43 | 10253,09 | 2565,4513 | 1946,186502 | CD630_01060 | cwlD E |
| CD630_10310 | 1687,181 | -2,311032 | 0,361489 | -6,393092 | 1,63E-10 | 7,46E-09 | CD630_10 | 2754,958 | 2943,357 | 656,5036 | 393,902816 | CD630_10310 | cell wall anchored protein F |
| CD630_21440 | 593,6612 | -2,56057 | 0,401525 | -6,37711 | 1,80E-10 | 8,19E-09 | CD630_21 | 988,8671 | 1073,71 | 210,75946 | 101,3083385 | CD630_21440 | membrane protein E/K |
| CD630_10633 | 3616,416 | -2,19056 | 0,344507 | -6,358538 | 2,04E-10 | 9,13E-09 | CD630_10 | 6250,216 | 5779,086 | 1455,936 | 980,4247756 | CD630_10633 | hypothetical protein E/K (operon) |
| CD630_01960 | 3211,861 | -2,070255 | 0,325802 | -6,354327 | 2,09E-10 | 9,28E-09 | CD630_01 | 4873,187 | 5635,366 | 1145,8531 | 1193,038986 | CD630_01960 | pseudogene K |
| CD630_19280 | 3801,19 | -2,91274 | 0,461114 | -6,316749 | 2,67E-10 | 1,17E-08 | CD630_19 | 5328,895 | 8394,401 | 1155,5432 | 325,9195889 | CD630_19280 | membrane protein E |
| CD630_10450 | 1540,333 | -2,181796 | 0,348792 | -6,255293 | 3,97E-10 | 1,72E-08 | CD630_10 | 3027,842 | 2093,928 | 525,68739 | 513,8732168 | CD630_10450 | sporulation integral membrane protein Spo0A |
| CD630_30230 | 979,3503 | -2,44551 | 0,39392 | -6,208131 | 5,36E-10 | 2,30E-08 | CD630_30 | 2223,6 | 1148,47 | 237,40721 | 307,9240288 | CD630_30230 | hypothetical protein |
| CD630_28450 | 6532,16 | -2,341367 | 0,377961 | -6,194737 | 5,84E-10 | 2,45E-08 | CD630_28 | 14642,08 | 7550,771 | 1756,3288 | 2179,462282 | CD630_28450 | Rbr rubrerythrin G |
| CD630_03140 | 3295,233 | -2,56584 | 0,417418 | -6,146931 | 7,90E-10 | 3,24E-08 | CD630_03 | 5027,191 | 6469,328 | 1218,5288 | 465,8850565 | CD630_03140 | membrane protein E |
| CD630_24010 | 7651,238 | -2,132895 | 0,347596 | -6,136131 | 8,46E-10 | 3,43E-08 | CD630_24 | 15297,72 | 9987,564 | 2463,7054 | 2855,962042 | CD630_24010 | CotD manganese catalase K |
| CD630_33120 | 1103,208 | -2,246171 | 0,366456 | -6,129435 | 8,82E-10 | 3,54E-08 | CD630_33 | 2179,47 | 1520,337 | 419,0964 | 293,927482 | CD630_33120 | major facilitator superfamily transporter G |
| CD630_21021 | 337,5277 | -2,893533 | 0,473368 | -6,112652 | 9,80E-10 | 3,90E-08 | CD630_21 | 575,4882 | 638,0389 | 106,59099 | 29,9926002 | CD630_21021 | hypothetical protein |
| CD630_26360 | 2631,326 | -2,489513 | 0,408507 | -6,094178 | 1,10E-09 | 4,33E-08 | CD630_26 | 5998,947 | 3114,145 | 491,77207 | 920,4395752 | CD630_26360 | membrane protein F/G |
| CD630_34620 | 14282,78 | -2,009038 | 0,329969 | -6,08856 | 1,14E-09 | 4,44E-08 | CD630_34 | 24927,92 | 21436,17 | 4731,1865 | 6035,844166 | CD630_34620 | mazE antitoxin endoAI E |
| CD630_11950 | 6191,056 | -2,286807 | 0,376664 | -6,07121 | 1,27E-09 | 4,85E-08 | CD630_11 | 8540,461 | 12354,11 | 2480,6631 | 1388,990641 | CD630_11950 | SpoIIIAD E |
| CD630_24890 | 1116,023 | -2,267613 | 0,373513 | -6,071047 | 1,27E-09 | 4,85E-08 | CD630_24 | 2247,016 | 1507,447 | 426,36396 | 283,2634464 | CD630_24890 | PTS operon transcription antiterminator |
| CD630_19981 | 291,6189 | -2,740359 | 0,451571 | -6,068498 | 1,29E-09 | 4,88E-08 | CD630_19 | 450,3038 | 582,6133 | 96,900901 | 36,65762247 | CD630_19981 | pseudogene |
| CD630_24740 | 883,3688 | -2,343502 | 0,388464 | -6,032735 | 1,61E-09 | 5,98E-08 | CD630_24 | 1850,748 | 1150,403 | 327,04054 | 205,2826858 | CD630_24740 | DNA polymerase III subunit delta |
| CD630_14630 | 13773,21 | -2,129003 | 0,35411 | -6,012266 | 1,83E-09 | 6,72E-08 | CD630_14 | 21974,82 | 23537,83 | 5995,7432 | 3584,448976 | CD630_14630 | hypothetical protein G |
| CD630_10630 | 3394,009 | -2,42459 | 0,404156 | -5,999139 | 1,98E-09 | 7,22E-08 | CD630_10 | 4324,717 | 7334,869 | 1283,9369 | 632,5106132 | CD630_10630 | hypothetical protein E/K (operon) |
| CD630_05720 | 7400,121 | -2,160309 | 0,3602 | -5,997519 | 2,00E-09 | 7,22E-08 | CD630_05 | 11068,47 | 13492,91 | 3166,2369 | 1872,871257 | CD630_05720 | YtxC Sporulation protein |
| CD630_15600 | 2551,136 | -2,500288 | 0,418397 | -5,975871 | 2,29E-09 | 8,12E-08 | CD630_15 | 3627,647 | 5215,807 | 973,85405 | 387,2377938 | CD630_15600 | hypothetical protein |
| CD630_06950 | 3564,209 | -2,205552 | 0,370463 | -5,953501 | 2,62E-09 | 9,12E-08 | CD630_06 | 5776,497 | 6127,751 | 1533,4568 | 819,1312367 | CD630_06950 | metalloendopeptidase |
| CD630_34920 | 5836,338 | -2,532283 | 0,426426 | -5,938391 | 2,88E-09 | 9,91E-08 | CD630_34 | 9581,563 | 10739,03 | 2252,9459 | 771,8095786 | CD630_34920 | DivIC septum formation initiation protein |
| CD630_07600 | 4372,152 | -2,129077 | 0,358748 | -5,93474 | 2,94E-09 | 1,00E-07 | CD630_07 | 7150,824 | 7299,423 | 1918,6378 | 1119,723741 | CD630_07600 | Ca2+/Na+ antiporter E |
| CD630_06750 | 321,0522 | -2,528265 | 0,431598 | -5,857913 | 4,69E-09 | 1,56E-07 | CD630_06 | 498,036 | 615,482 | 118,7036 | 51,98717369 | CD630_06750 | acetyltransferase |
| CD630_26350 | 3737,509 | -2,111593 | 0,361229 | -5,845585 | 5,05E-09 | 1,66E-07 | CD630_26 | 7247,189 | 5089,488 | 1034,4171 | 1578,943775 | CD630_26350 | Putative membrane protein G |
| CD630_34160 | 192,2695 | -2,402965 | 0,413603 | -5,809829 | 6,25E-09 | 2,01E-07 | CD630_34 | 310,7096 | 350,5991 | 48,45045 | 59,31869818 | CD630_34160 | sugar family ABC transporter permease |
| CD630_15180 | 1588,504 | -2,007914 | 0,346406 | -5,796426 | 6,77E-09 | 2,16E-07 | CD630_15 | 2446,05 | 2720,366 | 491,77207 | 695,8283248 | CD630_15180 | Ferrous iron transport protein FeoA K |
| CD630_34860 | 883,5554 | -2,002262 | 0,345655 | -5,792658 | 6,93E-09 | 2,19E-07 | CD630_34 | 1496,81 | 1370,172 | 370,64594 | 296,5934909 | CD630_34860 | membrane protein |
| CD630_36520 | 15804,15 | -2,25835 | 0,389932 | -5,791648 | 6,97E-09 | 2,19E-07 | CD630_36 | 21337,19 | 31903,23 | 6644,9793 | 3331,178129 | CD630_36520 | Putative peptidase M1 family E |
| CD630_16780 | 696,3766 | -2,029921 | 0,355789 | -5,705415 | 1,16E-08 | 3,52E-07 | CD630_16 | 1263,552 | 1010,55 | 230,13964 | 281,2639397 | CD630_16780 | membrane protein Spo0A |
| CD630_19050 | 1522,95 | -2,107289 | 0,370041 | -5,694741 | 1,24E-08 | 3,69E-07 | CD630_19 | 2071,397 | 2951,091 | 654,08108 | 415,2308873 | CD630_19050 | Site-specific recombinase (putative SpoIVCA homolog) Spo0A? |
| CD630_28000 | 3760,846 | -2,104723 | 0,370824 | -5,675792 | 1,38E-08 | 4,06E-07 | CD630_28 | 6626,67 | 5777,797 | 1715,1459 | 923,7720863 | CD630_28000 | Putative membrane protein E |
| CD630_12910 | 8596,751 | -2,13138 | 0,377259 | -5,649651 | 1,61E-08 | 4,59E-07 | CD630_12 | 17842,84 | 10643,65 | 2274,7486 | 3625,772114 | CD630_12910 | dacF G |
| CD630_28560 | 2047,518 | -2,301469 | 0,408679 | -5,631482 | 1,79E-08 | 4,92E-07 | CD630_28 | 3034,147 | 3906,216 | 881,7982 | 367,9092292 | CD630_28560 | membrane protein F |
| CD630_21210 | 1103,023 | -2,153435 | 0,383103 | -5,621038 | 1,90E-08 | 5,19E-07 | CD630_21 | 2297,45 | 1366,306 | 445,74414 | 302,592011 | CD630_21210 | hypothetical protein E |
| CD630_20141 | 379,6465 | -2,370396 | 0,422808 | -5,606314 | 2,07E-08 | 5,57E-07 | CD630_20 | 696,1696 | 598,7254 | 155,04144 | 68,64972936 | CD630_20141 | hypothetical protein |
| CD630_23160 | 204,8794 | -2,276539 | 0,406973 | -5,593835 | 2,22E-08 | 5,83E-07 | CD630_23 | 373,7521 | 320,3084 | 58,14054 | 67,3167249 | CD630_23160 | OmpR family E |
| CD630_13151 | 196,3057 | -2,646103 | 0,473659 | -5,586518 | 2,32E-08 | 6,04E-07 | CD630_13 | 380,957 | 309,3522 | 70,253153 | 24,66058239 | CD630_13151 | hypothetical protein |
| CD630_23740 | 7531,116 | -2,40867 | 0,431515 | -5,581889 | 2,38E-08 | 6,16E-07 | CD630_23 | 11761,93 | 14149,64 | 3163,8144 | 1049,074505 | CD630_23740 | hypothetical protein E |
| CD630_32970 | 4995,424 | -2,121845 | 0,380391 | -5,578066 | 2,43E-08 | 6,25E-07 | CD630_32 | 11072,07 | 5464,578 | 1652,1604 | 1792,89099 | CD630_32970 | hypothetical protein |
| CD630_11940 | 823,2558 | -2,144331 | 0,387142 | -5,538877 | 3,04E-08 | 7,48E-07 | CD630_11 | 949,2403 | 1785,864 | 271,32252 | 286,5959575 | CD630_11940 | SpoIIIAC E |
| CD630_28181 | 87,69277 | -2,848382 | 0,518344 | -5,495158 | 3,90E-08 | 9,41E-07 | CD630_28 | 174,7179 | 140,4975 | 24,225225 | 11,33053786 | CD630_28181 | hypothetical protein |
| CD630_10680 | 2888,899 | -2,353647 | 0,430447 | -5,467916 | 4,55E-08 | 1,08E-06 | CD630_10 | 4117,578 | 5758,462 | 1240,3315 | 439,2249674 | CD630_10680 | polysaccharide biosynthesis/spoVB E |
| CD630_28680 | 1808,09 | -2,200094 | 0,404825 | -5,43468 | 5,49E-08 | 1,28E-06 | CD630_28 | 4208,539 | 1849,024 | 658,92612 | 515,8727235 | CD630_28680 | oxidoreductase G |
| CD630_19410 | 7926,392 | -2,047055 | 0,378965 | -5,401706 | 6,60E-08 | 1,52E-06 | CD630_19 | 17047,6 | 8929,322 | 3275,2504 | 2453,394697 | CD630_19410 | hypothetical protein Spo0A |
| CD630_34391 | 4070,939 | -2,664584 | 0,494461 | -5,38887 | 7,09E-08 | 1,59E-06 | CD630_34 | 6685,21 | 7780,852 | 1523,7667 | 293,927482 | CD630_34391 | hypothetical protein |
| CD630_36361 | 173,0053 | -2,550241 | 0,473296 | -5,388254 | 7,11E-08 | 1,59E-06 | CD630_36 | 244,0646 | 358,9774 | 62,985585 | 25,99358684 | CD630_36361 | hypothetical protein E |
| CD630_11740 | 982,2937 | -2,001488 | 0,379064 | -5,280085 | 1,29E-07 | 2,72E-06 | CD630_11 | 1549,946 | 1644,722 | 477,23694 | 257,2698595 | CD630_11740 | hypothetical protein |
| CD630_12420 | 401,6451 | -2,164686 | 0,411514 | -5,260294 | 1,44E-07 | 2,98E-06 | CD630_12 | 759,2121 | 577,4574 | 179,26667 | 90,64430284 | CD630_12420 | hypothetical protein |
| CD630_19510 | 860,2831 | -2,088772 | 0,401005 | -5,208848 | 1,90E-07 | 3,83E-06 | CD630_19 | 1520,225 | 1316,036 | 414,25135 | 190,6196369 | CD630_19510 | acyl-CoA N-acyltransferase K |
| CD630_16840 | 702,296 | -2,04725 | 0,393577 | -5,201656 | 1,98E-07 | 3,93E-06 | CD630_16 | 947,4391 | 1353,416 | 327,04054 | 181,2886057 | CD630_16840 | radical SAM superfamily protein |
| CD630_20470 | 190,0058 | -2,396769 | 0,462286 | -5,184601 | 2,16E-07 | 4,28E-06 | CD630_20 | 244,0646 | 407,9582 | 72,675675 | 35,32461802 | CD630_20470 | OmpR family, |
| CD630_22640 | 394,4814 | -2,083905 | 0,402093 | -5,182647 | 2,19E-07 | 4,30E-06 | CD630_22 | 638,5307 | 659,3069 | 184,11171 | 95,97632066 | CD630_22640 | hypothetical protein |
| CD630_24680 | 1694,924 | -2,145206 | 0,420954 | -5,096061 | 3,47E-07 | 6,43E-06 | CD630_24 | 2468,565 | 3177,305 | 821,23513 | 312,5895444 | CD630_24680 | hypothetical protein F |
| CD630_11960 | 9204,655 | -2,281769 | 0,451057 | -5,058717 | 4,22E-07 | 7,61E-06 | CD630_11 | 10773,07 | 20535,19 | 4120,7108 | 1389,657143 | CD630_11960 | SpoIIIAE E |
| CD630_26600 | 10539,53 | -2,164738 | 0,431374 | -5,018241 | 5,21E-07 | 9,27E-06 | CD630_26 | 26401,31 | 8875,185 | 3030,5757 | 3851,049866 | CD630_26600 | M16 family peptidase F |
| CD630_33670 | 81,81567 | -2,493338 | 0,497122 | -5,015549 | 5,29E-07 | 9,31E-06 | CD630_33 | 149,5008 | 136,6306 | 21,802703 | 19,32856458 | CD630_33670 | hypothetical protein |
| CD630_00511 | 752,3756 | -2,05614 | 0,410131 | -5,013377 | 5,35E-07 | 9,38E-06 | CD630_00 | 856,4778 | 1617,01 | 336,73063 | 199,2841658 | CD630_00511 | hypothetical protein |
| CD630_21500 | 145,7368 | -2,207258 | 0,444601 | -4,964576 | 6,89E-07 | 1,16E-05 | CD630_21 | 198,1337 | 291,9511 | 50,872973 | 41,98964029 | CD630_21500 | endonuclease/exonuclease/phosphatase |
| CD630_24780 | 1160,398 | -2,039404 | 0,419388 | -4,862807 | 1,16E-06 | 1,88E-05 | CD630_24 | 2503,689 | 1308,302 | 552,33513 | 277,2649263 | CD630_24780 | DNA uptake transporter |
| CD630_14230 | 157,9891 | -2,01645 | 0,419794 | -4,803428 | 1,56E-06 | 2,44E-05 | CD630_14 | 250,3689 | 267,4607 | 58,14054 | 55,98618705 | CD630_14230 | hypothetical protein Spo0A |
| CD630_21860 | 377,7067 | -2,564003 | 0,533698 | -4,80422 | 1,55E-06 | 2,44E-05 | CD630_21 | 399,8697 | 933,8569 | 147,77387 | 29,32609798 | CD630_21860 | hypothetical protein |
| CD630_34540 | 367,6653 | -2,019504 | 0,44543 | -4,533831 | 5,79E-06 | 7,60E-05 | CD630_34 | 602,5064 | 601,9478 | 196,22432 | 69,98273381 | CD630_34540 | nucleotide-binding protein |
| CD630_25630 | 884,6557 | -2,18775 | 0,484507 | -4,515415 | 6,32E-06 | 8,26E-05 | CD630_25 | 2283,04 | 703,7762 | 385,18108 | 166,6255567 | CD630_25630 | hypothetical protein |
| CD630_28390 | 268,3914 | -2,004234 | 0,453314 | -4,421291 | 9,81E-06 | 0,000121 | CD630_28 | 392,6649 | 484,6518 | 142,92883 | 53,32017814 | CD630_28390 | two-component sensor histidine kinase |
| CD630_26650 | 160,7633 | -2,04083 | 0,462374 | -4,41382 | 1,02E-05 | 0,000124 | CD630_26 | 214,3446 | 313,8636 | 77,520721 | 37,3241247 | CD630_26650 | AraC family transcriptional regulator |
| CD630_17060 | 56,56029 | -2,512736 | 0,571183 | -4,399175 | 1,09E-05 | 0,000131 | CD630_17 | 82,85589 | 116,0071 | 19,38018 | 7,998026721 | CD630_17060 | thiamine-phosphate pyrophosphorylase ThiE2 |
| CD630_25970 | 73,80919 | -2,23814 | 0,511135 | -4,378773 | 1,19E-05 | 0,000142 | CD630_25 | 147,6996 | 104,4064 | 21,802703 | 21,32807126 | CD630_25970 | lspA lipoprotein signal peptidase |
| CD630_19260 | 65,14142 | -2,384046 | 0,561565 | -4,245364 | 2,18E-05 | 0,000241 | CD630_19 | 100,868 | 124,3854 | 26,647748 | 8,664528948 | CD630_19260 | pseudogene E |
| CD630_26870 | 1482,986 | -2,059974 | 0,491466 | -4,191489 | 2,77E-05 | 0,000298 | CD630_26 | 2486,577 | 2445,816 | 830,92522 | 168,6250634 | CD630_26870 | hypothetical protein F |
| CD630_02921 | 60,9862 | -2,344477 | 0,563145 | -4,163186 | 3,14E-05 | 0,000328 | CD630_02 | 123,3832 | 87,0053 | 24,225225 | 9,331031175 | CD630_02921 | hypothetical protein |
| CD630_33690 | 186,2889 | -2,118652 | 0,510064 | -4,153699 | 3,27E-05 | 0,000339 | CD630_33 | 236,8598 | 385,4013 | 96,900901 | 25,99358684 | CD630_33690 | hypothetical protein |
| CD630_07380 | 156,1926 | -2,08857 | 0,510141 | -4,094104 | 4,24E-05 | 0,000421 | CD630_07 | 183,7239 | 335,776 | 79,943243 | 25,32708462 | CD630_07380 | hypothetical protein |
| CD630_18210 | 55,73994 | -2,246545 | 0,562055 | -3,997023 | 6,41E-05 | 0,00059 | CD630_18 | 78,35285 | 112,1402 | 21,802703 | 10,66403563 | CD630_18210 | transposase-like protein B |
| EBG00001175819 | 42,12402 | -2,37458 | 0,603913 | -3,931986 | 8,42E-05 | 0,000744 | EBG00001 | 87,35893 | 59,93699 | 14,535135 | 6,665022268 | EBG00001175819 | |
| CD630_01631 | 46,06692 | -2,225907 | 0,588832 | -3,780206 | 0,000157 | 0,001259 | CD630_01 | 63,04253 | 97,96153 | 7,2675675 | 15,99605344 | CD630_01631 | hypothetical protein |
| CD630_15640 | 45,22533 | -2,035705 | 0,573016 | -3,552615 | 0,000381 | 0,002612 | CD630_15 | 62,14192 | 90,22772 | 14,535135 | 13,99654676 | CD630_15640 | hypothetical protein |
| CD630_17452 | 32,17147 | -2,263394 | 0,638756 | -3,543443 | 0,000395 | 0,002672 | CD630_17 | 55,83767 | 56,07008 | 12,112613 | 4,665515587 | CD630_17452 | hypothetical protein |
| EBG00001175749 | 43,31341 | -2,166945 | 0,617509 | -3,509173 | 0,00045 | 0,002946 | EBG00001 | 53,13584 | 94,73911 | 19,38018 | 5,998520041 | EBG00001175749 | |
| EBG00001175906 | 51,95933 | -2,075558 | 0,616338 | -3,367564 | 0,000758 | 0,004556 | EBG00001 | 58,53949 | 116,6516 | 26,647748 | 5,998520041 | EBG00001175906 | |
| CD630_23030 | 19,2831 | -2,34322 | 0,709018 | -3,304879 | 0,00095 | 0,005472 | CD630_23 | 28,81944 | 41,89144 | 2,4225225 | 3,999013361 | CD630_23030 | pseudogene |
| CD630_12341 | 28,37011 | -2,111583 | 0,678424 | -3,112483 | 0,001855 | 0,009541 | CD630_12 | 29,72005 | 68,31528 | 12,112613 | 3,332511134 | CD630_12341 | hypothetical protein (skin) |

| GeneID | Base mean | log2 (FC) | StdErr | Wald-Stats | p-value | p-adj | Gene ID | Counts-y1 | Counts-y2 | Counts-wt1 | Counts-wt2 | GeneID | |
|---|---|---|---|---|---|---|---|---|---|---|---|---|---|
| CD630_20541 | 3388,689 | 2,4491154 | 0,328855 | 7,447397 | 9,52E-14 | 9,42E-12 | CD630_205 | 985,2646 | 973,1704 | 5167,241 | 6429,08 | CD630_205 | hypothetical protein |
| EBG0000117568 | 961,005 | 2,7381324 | 0,386394 | 7,086382 | 1,38E-12 | 9,72E-11 | EBG000011 | 284,592 | 162,4099 | 2010,694 | 1386,325 | EBG00001175680 | |
| CD630_33790 | 495,9061 | 2,3011964 | 0,359443 | 6,402122 | 1,53E-10 | 7,12E-09 | CD630_337 | 143,1966 | 165,6323 | 922,9811 | 751,8145 | CD630_337 | conjugative transposon protein |
| CD630_33830 | 950,2159 | 2,1386087 | 0,34484 | 6,201741 | 5,58E-10 | 2,37E-08 | CD630_338 | 364,746 | 294,5291 | 1700,611 | 1440,978 | CD630_338 | ATPase |
| CD630_06052 | 5175,903 | 2,0030469 | 0,335211 | 5,975476 | 2,29E-09 | 8,12E-08 | CD630_060 | 2224,501 | 1684,68 | 9098,995 | 7695,435 | CD630_060 | hypothetical protein |
| EBG0000117567 | 719268,1 | 2,0929505 | 0,352718 | 5,933782 | 2,96E-09 | 1,00E-07 | EBG000011 | 274987,9 | 236343,8 | 1468729 | 897011,4 | EBG00001175675 | |
| CD630_00370 | 820,193 | 2,0215244 | 0,345894 | 5,844355 | 5,09E-09 | 1,66E-07 | CD630_003 | 267,4804 | 342,2209 | 1368,725 | 1302,345 | CD630_003 | acetoin dehydrogenase E1 component subunit beta |
| CD630_33911 | 1006,281 | 2,0876129 | 0,369923 | 5,643375 | 1,67E-08 | 4,73E-07 | CD630_339 | 368,3485 | 342,8653 | 2088,214 | 1225,698 | CD630_339 | hypothetical protein |
| CD630_33800 | 329,6459 | 2,1228161 | 0,378394 | 5,610072 | 2,02E-08 | 5,49E-07 | CD630_338 | 120,6814 | 106,3398 | 625,0108 | 466,5516 | CD630_338 | cell wall hydrolase |
| CD630_28720 | 185,5775 | 2,2579327 | 0,405231 | 5,57196 | 2,52E-08 | 6,43E-07 | CD630_287 | 53,13584 | 62,51492 | 336,7306 | 289,9285 | CD630_287 | uxaA-D-galactate dehydratase/altronate hydrolase |
| CD630_11031 | 562,9563 | 2,1024641 | 0,380489 | 5,525683 | 3,28E-08 | 7,96E-07 | CD630_110 | 169,3142 | 222,3469 | 1141,008 | 719,1559 | CD630_110 | Transposon protein |
| CD630_01721 | 78572,74 | 2,0393905 | 0,37383 | 5,455399 | 4,89E-08 | 1,16E-06 | CD630_017 | 37355,4 | 19806,27 | 144699,7 | 112429,6 | CD630_017 | ferredoxin |
| CD630_33810 | 774,1556 | 2,0296647 | 0,373064 | 5,440521 | 5,31E-08 | 1,25E-06 | CD630_338 | 261,1762 | 304,1963 | 1579,485 | 951,7652 | CD630_338 | pseudogene |
| CD630_31001 | 297,999 | 2,1744729 | 0,403365 | 5,390838 | 7,01E-08 | 1,59E-06 | CD630_310 | 111,6753 | 84,42737 | 615,3207 | 380,5728 | CD630_310 | hypothetical protein |
| CD630_26000 | 5442,931 | 2,2008025 | 0,416131 | 5,28872 | 1,23E-07 | 2,62E-06 | CD630_260 | 2106,521 | 1402,397 | 5278,677 | 12984,13 | CD630_260 | cstA Carbon starvation **IIID** |
| CD630_14150 | 246,4029 | 2,7815404 | 0,537831 | 5,171777 | 2,32E-07 | 4,52E-06 | CD630_141 | 40,52734 | 54,13663 | 750,982 | 139,9655 | CD630_141 | membrane protein |
| CD630_33350 | 146,735 | 2,1940253 | 0,458296 | 4,787356 | 1,69E-06 | 2,61E-05 | CD630_333 | 33,32248 | 58,00354 | 307,6604 | 187,9536 | CD630_333 | conjugative transposon protein |
| CD630_28700 | 103,0117 | 2,1143094 | 0,45109 | 4,687115 | 2,77E-06 | 4,03E-05 | CD630_287 | 32,42187 | 36,09109 | 169,5766 | 173,9571 | CD630_287 | 2-keto-3-deoxygluconate permease |
| CD630_33780 | 52,93474 | 2,3099151 | 0,566148 | 4,080057 | 4,50E-05 | 0,00044 | CD630_337 | 9,006075 | 18,04554 | 118,7036 | 65,98372 | CD630_337 | conjugative transposon protein |
| CD630_03700 | 58,67486 | 2,0357824 | 0,523691 | 3,887375 | 0,000101 | 0,000857 | CD630_037 | 22,51519 | 16,75658 | 84,78829 | 110,6394 | CD630_037 | HTH-type transcriptional regulator |
| CD630_20451 | 55,0322 | 2,2873115 | 0,646369 | 3,538706 | 0,000402 | 0,002697 | CD630_204 | 5,403645 | 19,33451 | 164,7315 | 30,6591 | CD630_204 | hypothetical protein |
