## Supplementary figures and images for "A regulatory protein that represses sporulation in *Clostridioides difficile*"

### Supplemental Figures

Martins *et al.* – Figure S1

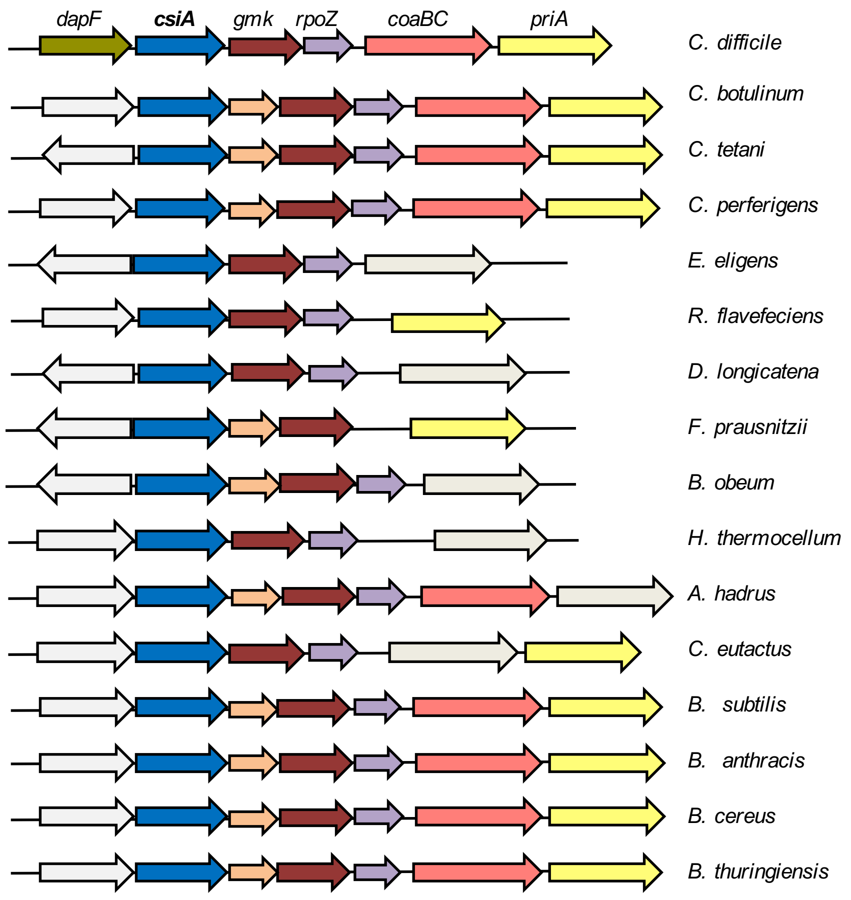

Martins *et al.* – Figure S2

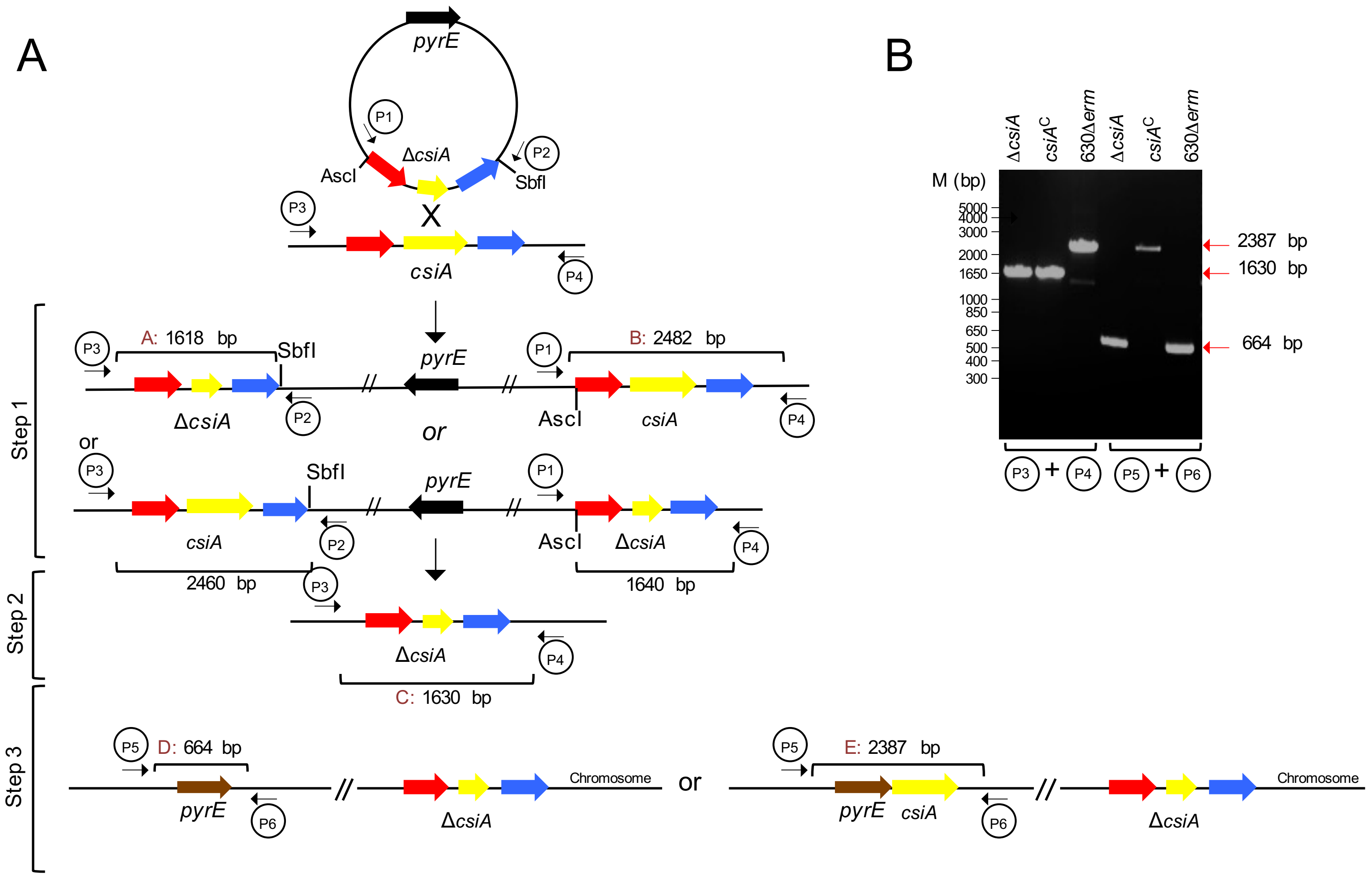

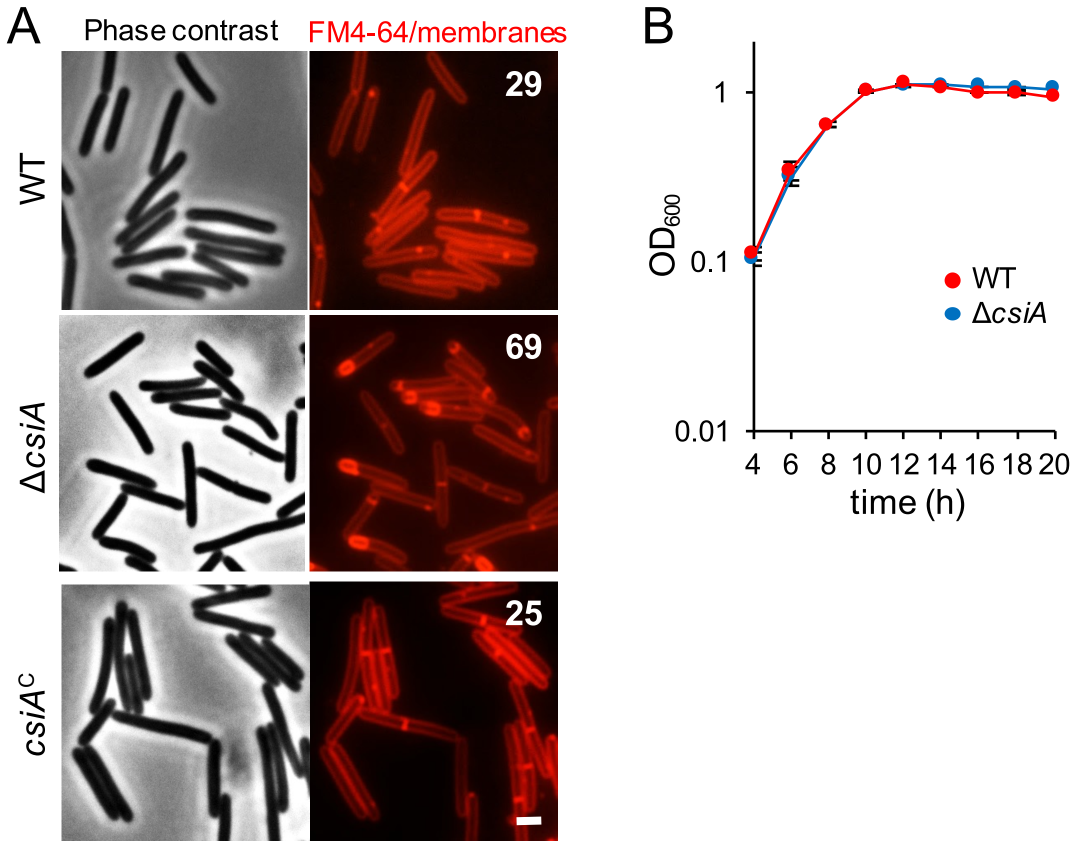

Martins *et al.* – Figure S4

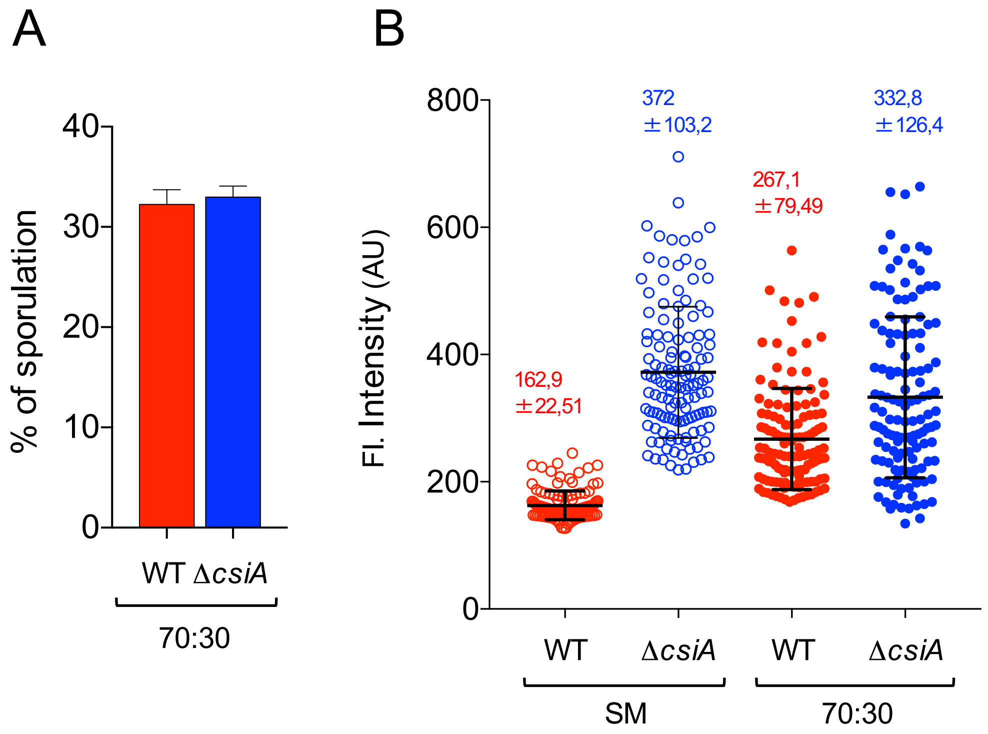

Martins *et al.* – Figure S5

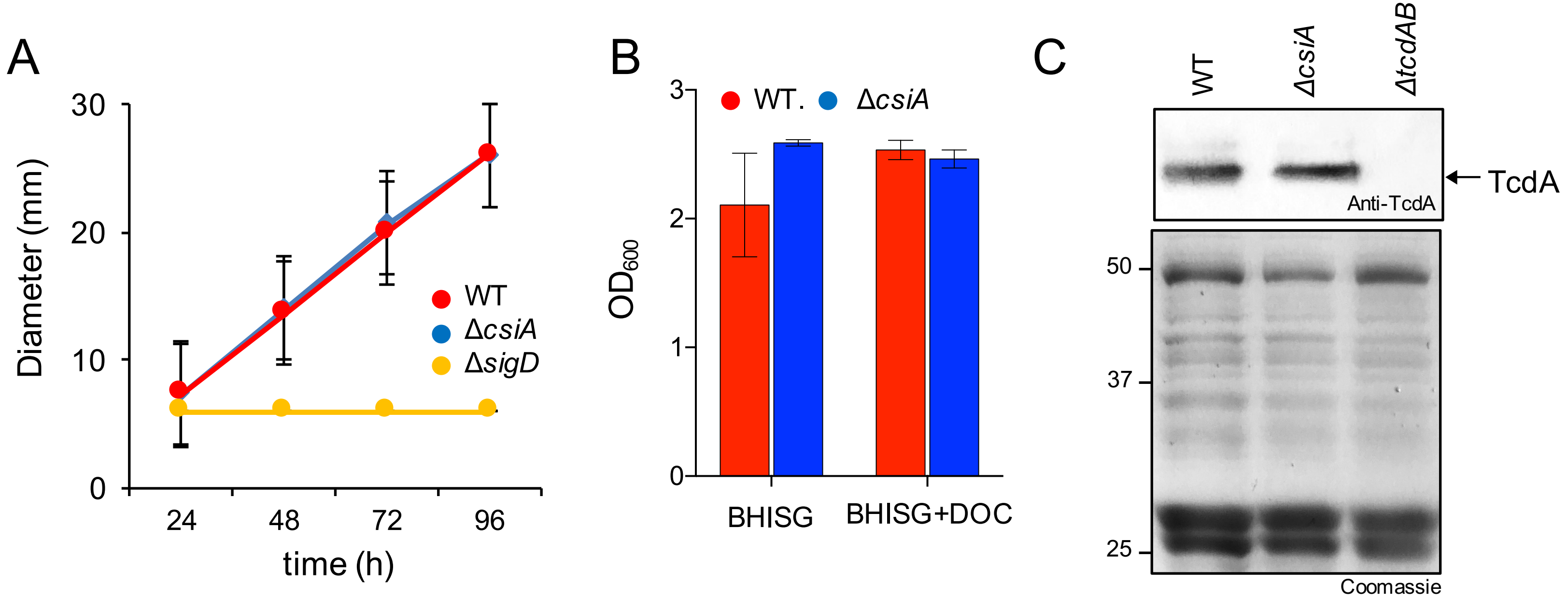
